# *De novo* indol-3-ylmethyl glucosinolate biosynthesis, and not long-distance transport, contributes to defence of Arabidopsis against powdery mildew

**DOI:** 10.1101/820720

**Authors:** Pascal Hunziker, Hassan Ghareeb, Lena Wagenknecht, Christoph Crocoll, Barbara Ann Halkier, Volker Lipka, Alexander Schulz

**Affiliations:** DynaMo Center, Department of Plant and Environmental Sciences, University of Copenhagen, Thorvaldsensvej 40, DK-1871 Frederiksberg C, Denmark; Department of Plant Cell Biology, Albrecht-von-Haller Institute of Plant Sciences, University of Goettingen, Julia-Lermontowa-Weg 3, D-37077 Göttingen, Germany; Central Microscopy Facility of the Faculty of Biology and Psychology, University of Goettingen, D-37077 Goettingen, Germany; Department of Plant Cell Biology, Goettingen Center for Molecular Biosciences (GZMB), University of Goettingen, D-37077 Goettingen, Germany

**Keywords:** Glucosinolate, biosynthesis, transport, powdery mildew, Arabidopsis, epidermis

## Abstract

Powdery mildew is a fungal disease that affects a wide range of plants and reduces crop yield worldwide. As obligate biotrophs, powdery mildew fungi manipulate living host cells to suppress defence responses and to obtain nutrients. Members of the plant order Brassicales produce indole glucosinolates that effectively protect them from attack by non-adapted fungi. Indol-3-ylmethyl glucosinolates are constitutively produced in the phloem and transported to epidermal cells for storage. Upon attack, indol-3-ylmethyl glucosinolates are activated by CYP81F2 to provide broad-spectrum defence against fungi. How *de novo* biosynthesis and transport contribute to defence of powdery mildew-attacked epidermal cells is unknown. Bioassays and glucosinolate analysis indicate that GTR glucosinolate transporters are not involved in antifungal defence. Using quantitative live-cell imaging of fluorophore-tagged markers, we show that accumulation of the glucosinolate biosynthetic enzymes CYP83B1 and SUR1 is induced in epidermal cells attacked by the non-adapted barley powdery mildew *Blumeria graminis* f.sp. *hordei.* By contrast, glucosinolate biosynthesis is attenuated during interaction with the virulent powdery mildew *Golovinomyces orontii*. Interestingly, SUR1 induction is delayed during the *Golovinomyces orontii* interaction. We conclude that epidermal *de novo* synthesis of indol-3-ylmethyl glucosinolate contributes to CYP81F2-mediated broad-spectrum antifungal resistance and that adapted powdery mildews may target this process.

---

Glucosinolates (GLS) are sulphur- and nitrogen-containing β-thioglycosides that are characteristic of the Brassicales order and function in defence against herbivores and pathogens. GLS are hydrolysed by myrosinases yielding toxic catabolites such as isothiocyanates or nitriles (Grubb & Abel, 2006). GLS hydrolysis pathways that function in herbivore defence rely on the destruction of cells and subsequent passive mixture of compartmentalized GLS and myrosinase (Halkier & Gershenzon, 2006). An additional pathway that operates cell-autonomously in living cells mediates broad-spectrum antifungal defence (Pawel Bednarek et al., 2009; Lipka et al., 2005). In *Arabidopsis thaliana* (hereafter Arabidopsis), this pathway depends on the myrosinase PEN2 (Lipka *et al*., 2005). PEN2-mediated indole GLS hydrolysis yields catabolites different from those detected upon wounding or herbivory (Agerbirk, De Vos, Kim, & Jander, 2009; Pawel Bednarek et al., 2009; Paweł Bednarek et al., 2011; Burow et al., 2008; Kim, Lee, Schroeder, & Jander, 2008; Piślewska-Bednarek et al., 2018). The PEN2-dependent defence pathway has been shown to restrict growth of the non-adapted biotrophic pathogens *Blumeria graminis* f. sp. *hordei*, *Erysiphe pisi* and hemibiotrophic *Phytophthora infestans*, the adapted biotrophic pathogens *Golovinomyces orontii* and *Golovinomyces cichoracearum* and the necrotrophic fungi *Plectosphaerella cucumerina* and *Botrytis cinerea* (Pawel Bednarek et al., 2009; Lipka et al., 2005; J. Xu et al., 2016). *PEN2* is constitutively expressed in epidermal cells and localized to peroxisomes and mitochondria where it is anchored to the membrane facing the cytosolic side (Fuchs *et al*., 2016; Lipka *et al*., 2005). Detailed studies revealed that mitochondrion-localized, but not peroxisome-localized PEN2 is required for non-host resistance towards *B. graminis* (Fuchs *et al*., 2016). Upon attack, PEN2-positive mitochondria are immobilized at the sites of attempted penetration (Fuchs *et al*., 2016). Moreover, the ER-anchored cytochrome P450 monooxygenase CYP81F2, which is not detectable in unchallenged epidermal cells, is cell-autonomously induced and reveals focal accumulation at *B. graminis* penetration sites, where it co-localizes with PEN2 (Fuchs *et al*., 2016). CYP81F2 catalyzes 4-hydroxylation of indol-3-ylmethyl GLS (I3M) and is required for synthesis of 4-methoxy-indol-3-ylmethyl GLS (4MOI3M), which is the relevant PEN2 substrate for production of antifungal compounds that effectively establish penetration resistance towards non-adapted powdery mildews (Pawel Bednarek et al., 2009; Hematy et al., 2019; Matern et al., 2019). I3M is the parent GLS of all modified indole GLS and produced from the precursor amino acid Trp via the GLS core structure synthesis pathway (Sønderby, Geu-Flores, & Halkier, 2010).

In the Arabidopsis ecotype Col-0, GLS core structure synthesis can be divided into two sub-pathways responsible for synthesis of Met-derived aliphatic and Trp-derived indole GLS, respectively (Sønderby *et al*., 2010). Core structure synthesis of indole GLS starts with the conversion of Trp to indole-3-acetaldoxime catalysed by CYP79B2 and CYP79B3, which function redundantly (Sønderby *et al*., 2010). The aldoxime is then further processed by CYP83B1/SUR2, GSTFs, GGP1, SUR1, UGT74B1 and SOTs to produce I3M (Sønderby *et al*., 2010). In contrast to SUR1, which is the only enzyme that converts *S*-alkyl-thiohydroximate into thiohydroximate and required for synthesis of both aliphatic and indole GLS, CYP83B1 is specific for indole GLS core structure synthesis. The equivalent reaction in the aliphatic sub-pathway is catalysed by CYP83A1. The two homologous CYP83s are both non-redundant and are therefore used as markers for the two sub-pathways (Naur et al., 2003; Nintemann et al., 2018). Both CYP83s are predominantly localized to the vasculature under normal growth conditions. Cellular localization in flower stalks revealed that CYP83B1 is exclusively localized to the phloem, while CYP83A1 was additionally found in the starch sheath and in xylem parenchyma (Nintemann *et al*., 2018). However, the localization of GLS synthesis in leaves has not been demonstrated on the cellular level yet. Presence of GLS synthesis in the vasculature as well as absence in the mesophyll and epidermis has been indicated by untargeted proteomics of dissected leaves (Svozil, Gruissem, & Baerenfaller, 2015). Despite the absence of GLS biosynthesis, high concentrations of GLS, particularly I3M, have been detected in the epidermis of leaves (O A Koroleva et al., 2000; Olga A Koroleva, Gibson, Cramer, & Stain, 2010; Madsen, Olsen, Nour-Eldin, & Halkier, 2014), indicating that GLS are transported from the site of synthesis (i.e. the vasculature) to the site of storage (i.e. epidermis).

To date, three plasma membrane-localized GLS transporters have been identified. GTR1/NPF2.10 and GTR2/NPF2.11 show proton-coupled import of both aliphatic and indole GLS into *Xenopus laevis* oocytes (Nour-Eldin *et al*., 2012). By contrast, the recently identified GTR3/NPF2.9 shows high specificity for indole GLS (Jørgensen *et al*., 2017). The current model suggests that GTRs affect seed loading, root exudation, intra-leaf distribution and transport of GLS between root, shoot and flower stalks via phloem loading and xylem retrieval (Andersen & Halkier, 2014; Andersen et al., 2013; Jørgensen et al., 2017; Madsen, Kunert, Reichelt, Gershenzon, & Halkier, 2015; Madsen et al., 2014; Nour-Eldin et al., 2012; D. Xu et al., 2016). This idea is supported by the vascular localization of GTR1-3 (Nour-Eldin et al., 2012; Wang & Tsay, 2011). In addition, cell-to-cell transport of GLS has been proposed to follow the symplasmic route by diffusion through plasmodesmata (Andersen et al., 2013; Hunziker, Halkier, & Schulz, 2019; Madsen et al., 2014; Nintemann et al., 2018; D. Xu et al., 2016). While side chain modifications of I3M and subsequent hydrolysis upon pathogen attack are well studied, relatively little is known about how the plant orchestrates I3M biosynthesis and transport to defend epidermal cells.

Here, we show that I3M is *de novo* synthesized in epidermal cells upon attack by powdery mildews. Qualitative and quantitative bioimaging revealed accumulation of fluorophore-tagged CYP83B1 in epidermal cells attacked by the powdery mildews *B. graminis* or *G. orontii*. However, accumulation of 4MOI3M was solely observed during the incompatible interaction with *B. graminis*. No increase of 4MOI3M was observed during the compatible interaction with *G. orontii*, despite successful induction of CYP81F2, indicating insufficient core-structure synthesis. Supporting this hypothesis, we show that induction of SUR1 is delayed in response to *G. orontii* compared to *B. graminis* suggesting that SUR1 is a potential *G. orontii* effector target. Moreover, we demonstrate that GTR-mediated GLS transport is not required for defence against powdery mildews, highlighting the importance of cell-autonomous defence biochemistry in plant immunity.

## Material and methods

### Plant growth and inoculations

*Arabidopsis thaliana* (L.) Heynh. plants were grown in a walk-in climate chamber under short-day conditions (8 h photoperiod, 22°C day, 18°C night, 65% relative humidity and 150 µmol m^-2^ s^-1^) for 4 weeks following vernalization at 4°C for 2 days. The Columbia-0 accession (Col-0) was used as wild-type. The following previously described T-DNA lines were used: *pen2-1* (Lipka *et al*., 2005), *cyp81F2-2* (Pawel Bednarek et al., 2009), *eds1-2* (Aarts *et al*., 1998), *edr1* (Frye & Innes, 1998), *gtr1gtr2* (Nour-Eldin *et al*., 2012) and *gtr1gtr2gtr3* (Jørgensen *et al*., 2017). The following previously described transgenic lines were used: *pCYP81F2::CYP81F2-RFP* in *cyp81F2-2* (Fuchs *et al*., 2016), *pGTR1::GTR1-YFP* in *gtr1gtr2* (Nour-Eldin *et al*., 2012), *pGTR2::GTR2-mOrange2* in *gtr1gtr2* (Nour-Eldin *et al*., 2012), *pCYP83A1::CYP83A1-mVenus* (D. Xu et al., 2016), *pCYP83B1::CYP83B1-mVenus* (D. Xu et al., 2016) and *pSUR1::SUR1-mVenus* (D. Xu et al., 2016). Positive T_2_ transformants either heterozygous or homozygous for the *pCYP83A1::CYP83A1-mVenus*, *pCYP83B1::CYP83B1-mVenus* or *pSUR1::SUR1-mVenus* transgenes were selected by germination on solid half-strength Murashige and Skoog medium containing 1% (w/v) Suc and 100 µg ml^-1^ hygromycin B. Seedlings were transferred to soil after 10 days. To produce conidiospores, *Blumeria graminis* f. sp. *hordei* isolate K1 was grown on *Hordeum vulgare* cv Ingrid (line l-10) for 10 to 14 d prior to inoculation. To produce conidiospores, *Golovinomyces orontii* was grown on the Arabidopsis accession Col-0 for 10 to 14 d prior to inoculation. Plants were randomized in trays and inoculated using a settling tower (Lipka *et al*., 2005). For *G. orontii* inoculations, the settling tower was equipped with a mesh screen. Plants were inoculated at noon (between 12.00 and 13.00 o’clock) and thereafter grown in reach-in growth chambers using the settings described in the previous paragraph.

### Bioassays

For *B. graminis* penetration resistance assays, fifth true leaves were harvested at 72 hpi and fixed in 80% ethanol, cleared for 10-14 d and subsequently subjected to aniline blue staining (150mM KH_2_PO_4_, pH 9.5 adjusted with KOH; 0.01% (w/v) aniline blue) overnight in the dark. Next, fungal structures were stained using an ethanolic solution of 0.6% Coomassie Brilliant Blue, washed in MiliQ-water and mounted in 50% glycerol. Samples were observed using a DM750 epifluorescence microscope with UV excitation and a long-pass UV filter set (Lipka *et al*., 2005). For sporulation bioassays with *G. orontii*, five-six plants were pooled at 12 dpi. 5 µL mg^-1^ MiliQ-water was added and spores were released by vortexing. 100 µL spore solution were mounted on a hemocytometer and counted using a DM750 microscope. Counts for each pool were technically replicated six times and the mean thereof was used as single biological replicate.

### Confocal microscopy and quantification of fluorescence

Fifth true leaves were harvested using forceps and mounted in a Calcofluor white solution (Fluorescent brightener 28). Samples were observed using a Leica TCS SP5 (Leica Microsystems, Wetzlar, Germany) equipped with argon and diode lasers. An HCX PL APO CS 20.0×0.70 DRY UV objective was used throughout the study. Scanning speed was set to 400 Hz in bidirectional mode. Zoom was set to 3.6 fold. Z-stack volume was set to 11 µm imaged in 1µm steps (11 steps/stack) starting on top of the adaxial epidermis with the conidiospore in focus. Sequential scanning between stacks was applied to reduce photobleaching. The first sequence was used to image fluorescent protein fusions using the argon laser with 20% pre-set power and a line average of 3. The 514 nm laser line with an AOBS setting of 12% was used for excitation of mVenus and mOrange2. mVenus emission light was recorded using a HyD hybrid detector with a detection window of 520-550 nm and a gain of 187 V. mOrange2 emission light was recorded using a HyD hybrid detector with a detection window of 520-575 nm and a gain of 187 V. Chlorophyll autofluorescence was recorded using a photomultiplier tube and a detection window of 681-732 nm and a gain of 561 V. The 561 nm laser line with an AOBS setting of 15% was used for excitation of RFP. RFP emission light was recorded using a HyD hybrid detector with a detection window of 580-620 nm and a gain of 277 V. Chlorophyll autofluorescence was recorded using a photomultiplier tube with a detection window of 678-731 nm and a gain of 553 V. The second sequence was used to image the Calcofluor white signal using the 405 nm diode UV laser with an AOBS setting of 4-5%. Calcofluor white emission light was recorded with a detection window of 420-460 nm and 98-296 V gain. The HyD detectors were used in standard acquisition mode. For quantification of mean fluorescence intensities, z-stacks were maximum projected and three lines were manually drawn across attacked cells to obtain three representative measurements. The mean of the three measurements was used for further analysis. ImageJ (version 2.0.0-rc-68/1.52i, http://imagej.net) was used for fluorescence quantification. Fixation and ClearSee treatment were performed as previously described.

### Desulfo-glucosinolate analysis by LC-MS

Entire fifth true leaves of 4-week-old plants were gently harvested by detachment at the proximal part of the petiole using fine forceps and immediately frozen in liquid nitrogen. Subsequently, samples were lyophilized for 24 hours, ground into fine powder and extracted in 85% methanol containing 50 µM *p*-hydroxybenzyl glucosinolate as internal standard as previously described (Andersen *et al*., 2013). Samples were 10-fold diluted with deionized water and subjected to analysis by LC-MS as previously described (Crocoll, Halkier, & Burow, 2016; Jensen, Jepsen, Halkier, Kliebenstein, & Burow, 2015).

### Statistical analysis

ANOVA analyses were performed in R version 3.4.2 (2017-09-28; https://www.R-project.org).

## Results

### GTRs are not involved in defence against *B. graminis* and *G. orontii*

To explore the role of transmembrane GLS transport in defence against adapted and non-adapted powdery mildews, we investigated whether accumulation of GTR1 and GTR2 – the *GTR*s expressed in aboveground tissue – is induced upon attack by non-adapted *B. graminis* or virulent *G. orontii*. As the hypothesized pathogen-induced expression of *GTR1* and/or *GTR2* is assumed to occur locally – if not cell-autonomously – and restricted to the epidermis, we used live-cell imaging of protein-fluorophore fusions instead of e.g. qualitative PCR that might not be able to detect perturbations in a small subpopulation of attacked epidermal cells when sampling whole leaves. We inoculated transgenic plants homozygous for *GTR1-YFP* or *GTR2-mOrange2* fluorophore fusions (*GTR1-YFP/gtr1gtr2* and *GTR2-mOrange2/gtr1gtr2*, respectively) and examined their expression at individual interaction sites on the fifth true leaf of 4-week-old plants. Both constructs are expressed under the control of their endogenous 5’ regulatory sequences in the *gtr1gtr2* double knockout background (Nour-Eldin *et al*., 2012). We imaged plants at 24 and 48 hours post inoculation (hpi) for several reasons: Firstly, pathogen induction of CYP81F2 has been reported to reach a maximum at 6 to 24 hpi with *B. graminis* and decrease to basal levels at 48 hpi (Fuchs *et al*., 2016). Secondly, 4MOI3M levels have been reported to be unaffected at 24 hpi with *G. orontii*, but showed two-fold induction at 48 hpi (Schön *et al*., 2013). Thirdly, the two time points allow to capture abundant attempted penetration events for both pathogens at 24 hpi and failure or success of penetration for *B. graminis* and *G. orontii*, respectively, at 48 hpi. The fluorescence signals of GTR1-YFP and GTR2-mOrange2 were close to the detection limit in unchallenged epidermal cells at both time points (Fig. 1), but showed prominent signals in the vasculature of unchallenged plants (Fig. S1). No pathogen-induced accumulation was observed at 24 hpi and 48 hpi with either *B. graminis* or *G. orontii* (Fig. 1), as also evident from quantitative analysis of mean fluorescence intensities (Fig. S2 and Fig. S3). Hence, accumulation of neither GTR1 nor GTR2 is induced upon attack by *B. graminis* or *G. orontii*.

**Figure 1:**
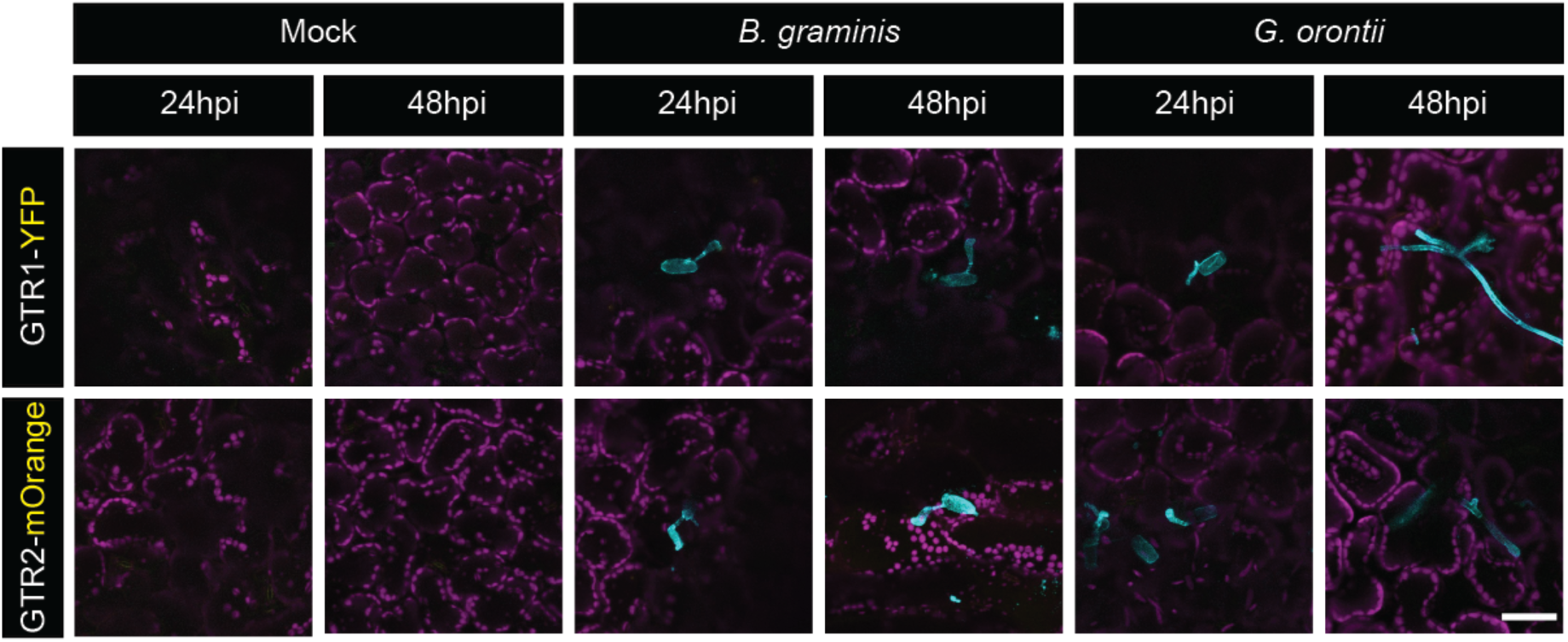
GTR1-YFP and GTR2-mOrange2 do not accumulate in attacked epidermal cells. Representative confocal micrographs of plants expressing *GTR1-YFP* or *GTR2-mOrange2* under the control of their native promoters at 24 hours post inoculation (hpi) and 48 hpi with *B. graminis* or *G. orontii* and following mock-treatment. All images represent z-projections through the adaxial epidermal cell layer. All images are overlays of YFP or mOrange2 fluorescence (yellow), chlorophyll autofluorescence (magenta) and calcofluor white staining (cyan). Scale bar = 50 µm.

To validate our results, we conducted quantitative bioassays using *B. graminis* and *G. orontii* on *gtr* knockout mutants by inoculating GLS transporter mutants with the virulent *G. orontii* and quantifying the number of spores produced at 12 days post inoculation (Fig. 2a). Loss-of-function mutants of *ENHANCED DISEASE RESISTANCE1 (EDR1)* and *ENHANCED DISEASE SUSCEPTIBILITY1 (EDS1)* significantly decrease and increase spore production of *G. orontii*, respectively (Aarts et al., 1998; Frye & Innes, 1998), and were therefore included as controls. As expected, spore production on *edr1* was significantly decreased while being increased on *eds1-2* compared to wild-type. On *pen2-1* mutants, *G. orontii* produced significantly more spores compared to wild-type and reached *eds1-2* levels. Hence, PEN2-mediated hydrolysis of GLS contributes to basal resistance against virulent *G. orontii*. When we examined the conidiospore production in double and triple mutants of the functionally redundant GTR1, GTR2 and GTR3 transporters (Jørgensen *et al*., 2017), we observed that the number of spores on *gtr1gtr2* and *gtr1gtr2gtr3* were not significantly different from wild-type. Compared to *pen2-1*, both double and triple *gtr* mutants showed significantly lower spore numbers. As GTR-mediated import of GLS from the apoplast would occur upstream of PEN2-mediated GLS hydrolysis, we expected a *pen2*-like phenotype and thus conclude that GTR1, GTR2 and GTR3 are not required for defence against *G. orontii*. Consistent with these findings, *GTR1-YFP/gtr1gtr2* (equivalent to *gtr2* single mutants) and *GTR2-mOrange2/gtr1gtr2* (equivalent to *gtr1* single mutants) showed no enhanced susceptibility towards *G. orontii*.

**Figure 2:**
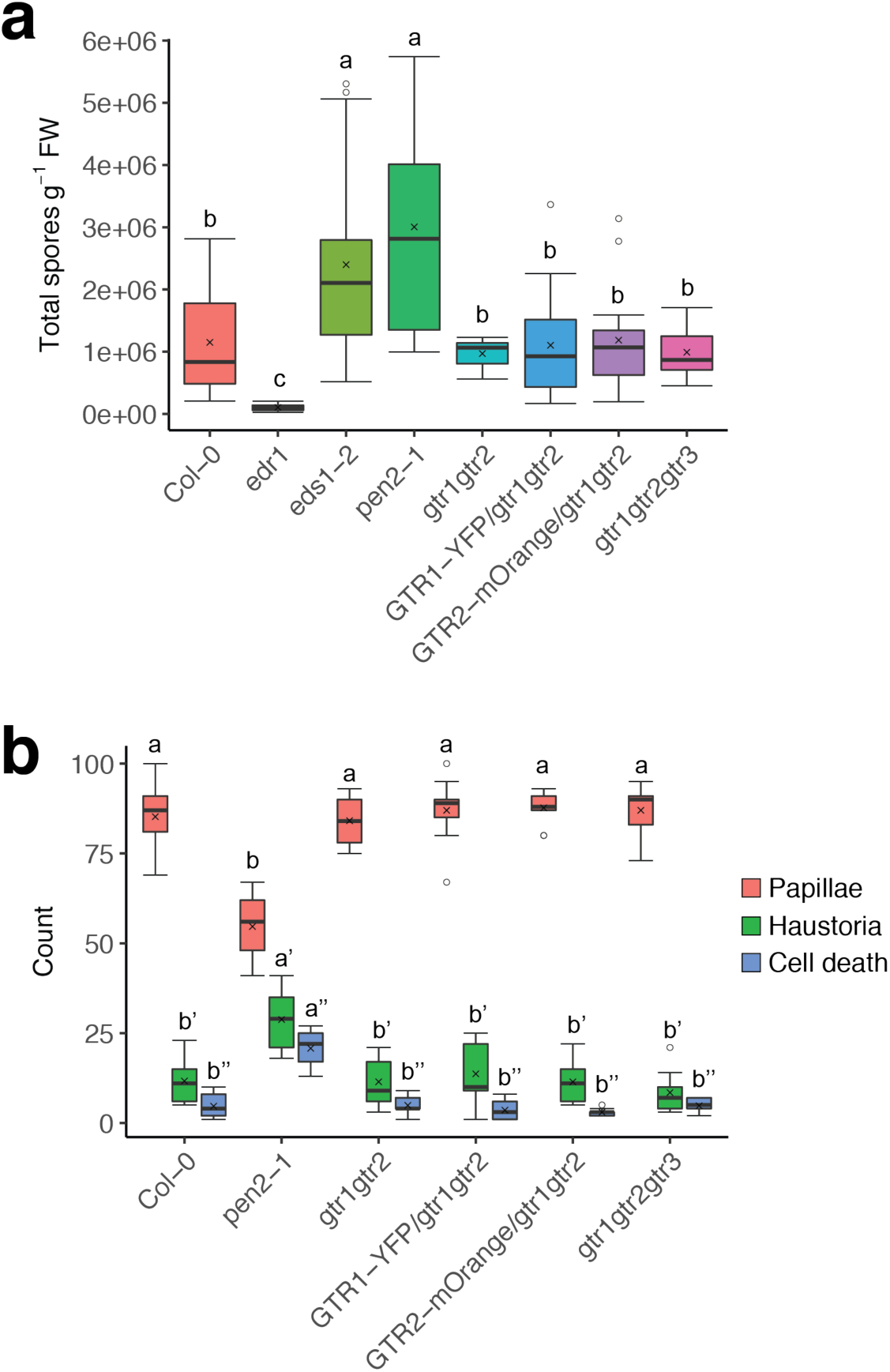
GTR1, GTR2 and GTR3 are not involved in defence against *B. graminis* and *G. orontii*. (a) Reproduction of *G. orontii* on wild-type (Col-0), *edr1* (negative control), *eds1-2* (positive control), *pen2-1*, *gtr1gtr2*, *GTR1-YFP/gtr1gtr2*, *GTR2-mOrange2/gtr1gtr2* and *gtr1gtr2gtr3* at 12 dpi. Spore numbers were determined for 4-6 pools of five plants. Pooled data from three independent experiments are shown. Individual boxplots show median (center line), mean (cross), first quartile (lower hinge), third quartile (upper hinge), whiskers (extending 1.5 times the inter-quartile range) and possible outliers (circles). Letters indicate significant differences between genotypes as determined by two-way ANOVA (p<0.001) with Tukey HSD post-hoc test. (b) Penetration resistance of wild-type (Col-0), *pen2-1*, *gtr1gtr2*, *GTR1-YFP/gtr1gtr2*, *GTR2-mOrange2/gtr1gtr2* and *gtr1gtr2gtr3* towards *B. graminis* at 72 hpi. Number of papillae, encased haustoria and cell death were scored for 100 interaction sites on the 5^th^ leaves of three individual plants per genotype. Pooled data from three independent experiments are shown. Letters indicate significant differences between genotypes as determined by two-way ANOVA (p<0.001; n = 3) with Tukey HSD post-hoc test.

When we challenged the double and triple *gtr* mutants with *B. graminis* and scored interaction sites at 72 hpi, the number of papillae, encased haustoria and epidermal cells undergoing cell death on *B. graminis*-inoculated *gtr1gtr2*, *GTR1-YFP/gtr1gtr2*, *GTR2-mOrange2/gtr1gtr2* and *gtr1gtr2gtr3* were not significantly different from wild-type at 72 hpi (Fig. 2b). Compared to wild-type and *gtr* mutants, *pen2-1* plants showed significantly more encased haustoria and cell death while the number of efficient papillae significantly decreased, indicating that more penetration attempts were successful. Moreover, we observed secondary hyphae at conidia with encased haustoria (both with and without cell death). These results suggest that GLS transporters are not required for penetration resistance towards *B. graminis*. In conclusion, GTR-mediated GLS transport is not involved in defence against *B. graminis* and *G. orontii*.

### *B. graminis* and *G. orontii* induce accumulation of key enzymes of indole GLS core structure synthesis in the epidermis

Assuming that epidermal cells challenged by powdery mildews do not receive GLS via transport, they must either remobilize preformed I3M intracellularly or synthesize it *de novo* via the core structure pathway. To test whether I3M is synthesized *de novo*, we inoculated transgenic plants expressing *pCYP83A1::CYP83A1-mVenus* or *pCYP83B1::CYP83B1-mVenus* with *B. graminis* and *G. orontii* and subsequently analyzed the accumulation of the fusion proteins using CLSM (Fig. 3). Plants expressing *pCYP81F2::CYP81F2-RFP* were included as a positive control for induction of I3M-to-4MOI3M conversion. The fluorescence signals of fusion proteins were below the detection limit in epidermal cells of unchallenged plants (Fig. 3a). However, the markers for core structure synthesis were detectable in vascular parenchyma cells of cleared leaves (Fig. S1) and transverse sections of non-cleared leaves (data not shown) of unchallenged plants. Upon inoculation with *B. graminis* or *G. orontii*, we observed prominent mVenus fluorescence in attacked epidermal cells of three independent *CYP83B1-mVenus*-expressing plant lines, while mVenus fluorescence in three independent *CYP83A1-mVenus* lines was below the detection limit (Fig. 3a). Quantification of mean fluorescence intensities in attacked cells revealed a statistically significant increase of the CYP83B1-mVenus signal at both 24 hpi and 48 hpi with both pathogens (Fig. 3b). Hence, both *B. graminis* and *G. orontii* induce the accumulation of key enzymes involved in indole GLS core structure synthesis in the epidermis.

**Figure 3:**
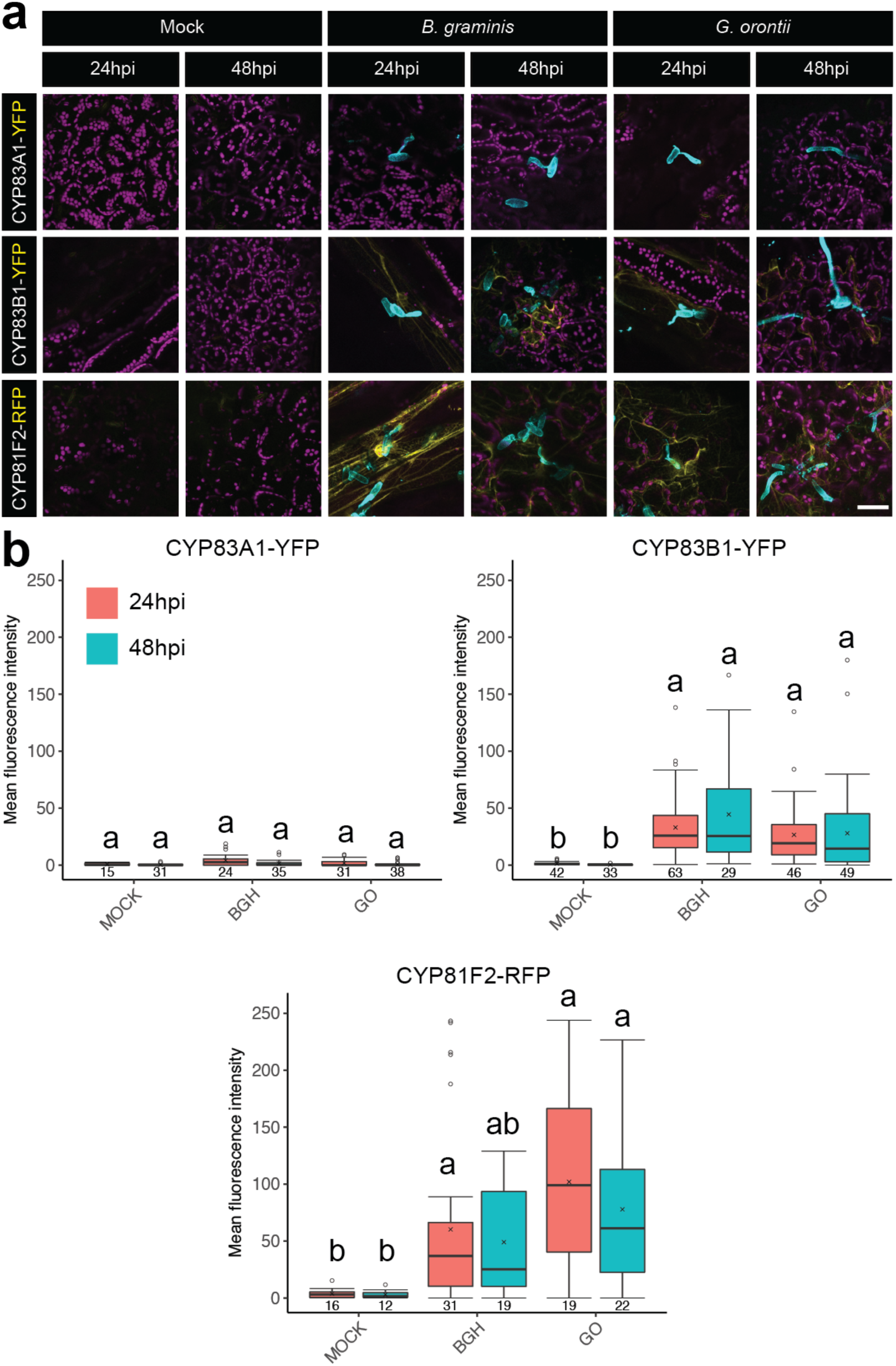
*B. graminis* and *G. orontii* induce markers for biosynthesis of indole glucosinolate core structure and side chain modification in attacked epidermal cells. (a) Representative confocal micrographs of transgenic plants expressing CYP83A1-YFP, CYP83B1-YFP and CYP81F2-RFP under the control of their native promoters at 24 hours post inoculation (hpi) and 48 hpi with *B. graminis* or *G. orontii* and following mock-treatment. All images represent z-projections through the adaxial epidermal cell layer. All images are overlays of YFP or RFP channels depicted in yellow, chlorophyll autofluorescence in magenta and calcofluor white staining in cyan. Scale bar = 50 µm. (b) Quantification of mean fluorescence intensities of CYP83A1-YFP, CYP83B1-YFP and CYP81F2-RFP following mock-treatment (MOCK) or inoculation with *B. graminis* (BGH) or *G. orontii* (GO) for 24 hpi (red boxes) and 48 hpi (blue boxes). Fluorescence was corrected by subtracting autofluorescence as determined in Col-0 plants (see Fig. S2). Results from three independent lines and three independent experiments were pooled for all genotypes except CYP81F2-RFP (only one line). Individual boxplots show median (center line), mean (cross), first quartile (lower hinge), third quartile (upper hinge), whiskers (extending 1.5 times the inter-quartile range) and possible outliers (circles). Values below each box indicate the number of observations. Letters indicate significant differences between treatment × timepoint interactions as determined by two-way ANOVA (p<0.05) with Tukey HSD post-hoc test.

### *B. graminis*, but not *G. orontii*, triggers 4MOI3M accumulation

To test whether both pathogens elicit the accumulation of GLS at the time points used in this study, we challenged wild-type plants with *B. graminis* or *G. orontii* and analyzed the GLS concentration in the fifth true leaf at 24 hpi and 48 hpi by LC-MS (Fig. 4 and S5). 4MOI3M levels were significantly induced in response to *B. graminis* at both time point (Fig. 4). By contrast, 4MOI3M levels were not significantly induced upon treatment with *G. orontii*. Hence, non-adapted *B. graminis*, but not virulent *G. orontii* triggers 4MOI3M accumulation. Total indole GLS did not reveal significant changes in response to the pathogen treatments, indicating a balance among individual indole GLS (Fig. 4). However, wild-type plants challenged by *G. orontii* revealed the overall highest indole GLS levels at 24 hpi and overall lowest levels at 48 hpi, reflecting a significant decrease of total indole GLS between 24 hpi and 48 hpi. This shift was reflected by I3M, which also showed a significant difference between 24 hpi and 48 hpi with *G. orontii*. Hence, *G. orontii* might trigger CYP81F2-catalyzed I3M-to-4MOI3M conversion and subsequent PEN2-mediated 4MOI3M hydrolysis, leading to a slight depletion of the I3M pool without significant accumulation of 4MOI3M. However, absence of complete I3M depletion indicates that most I3M is not accessible to ER-anchored CYP81F2 and mitochondria-anchored PEN2. Moreover, as induction of CYP81F2 accumulation is similar between *B. graminis* and *G. orontii* treatments (Fig. 3), we conclude that absence of 4MOI3M accumulation in response to *G. orontii* results from insufficient indole GLS core structure synthesis in the epidermis.

**Figure 4:**
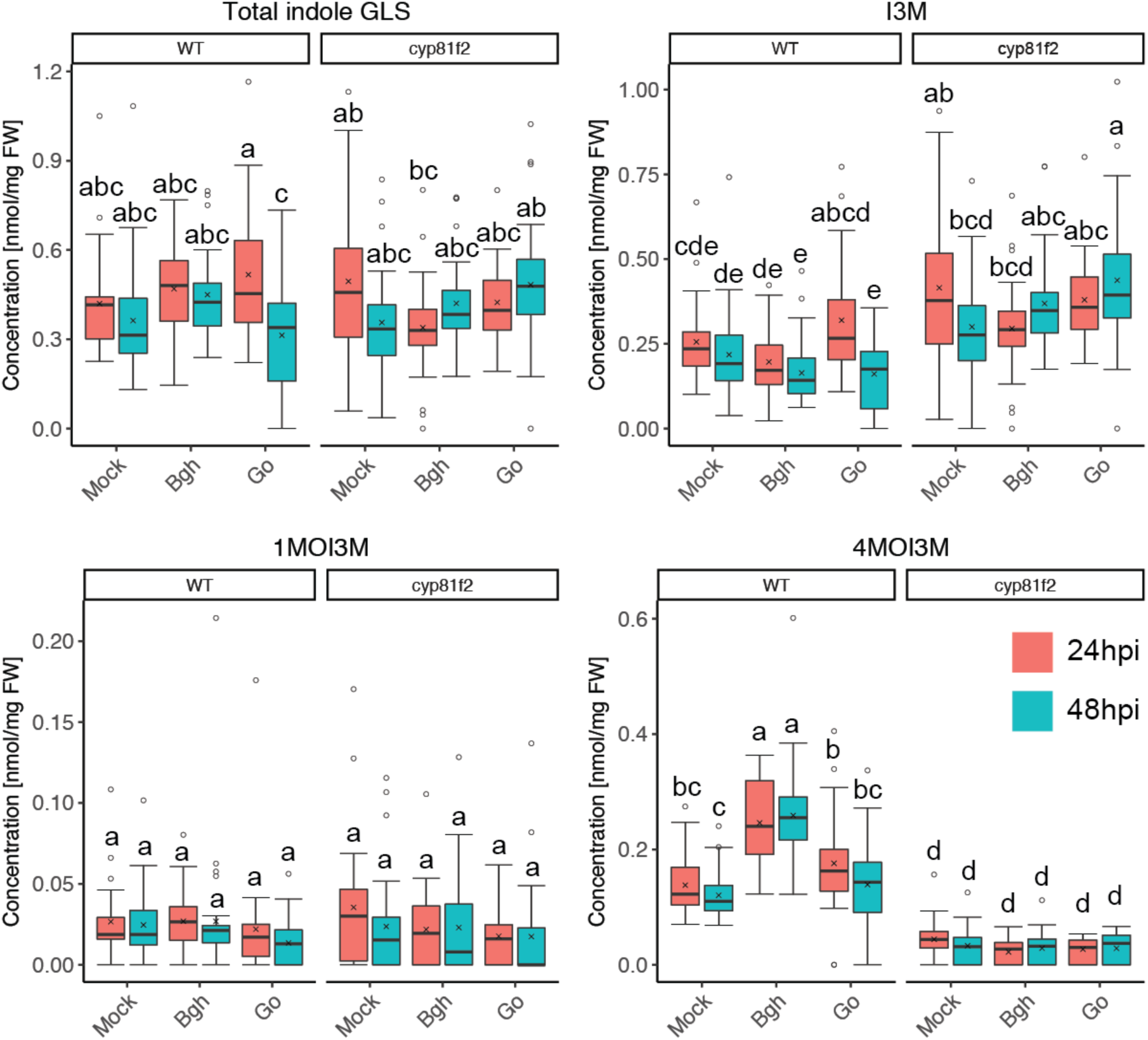
*B. graminis*, but not *G. orontii* triggers 4-methoxy-indol-3-ylmethyl glucosinolate accumulation. Quantification of total indole, indol-3-ylmethyl (I3M), 1-methoxy-indol-3-ylmethyl (1MOI3M) and 4-methoxy-indol-3-ylmethyl (4MOI3M) glucosinolates in whole leaves of Col-0 (wild-type; WT) and *cyp81f2* mutant plants following mock-treatment (Mock) or inoculation with *B. graminis* (Bgh) or *G. orontii* (Go) for 24 hours (red boxes) and 48 hours (blue boxes). Total indole glucosinolates represent the sum of I3M, 1MOI3M and 4MOI3M. Pooled data from three independent experiments are shown. Individual boxplots show median (center line), mean (cross), first quartile (lower hinge), third quartile (upper hinge), whiskers (extending 1.5 times the inter-quartile range) and possible outliers (circles). Letters indicate significant differences between treatment × timepoint interactions as determined by two-way ANOVA (p<0.05; n = 30) with Tukey HSD post-hoc test.

To gain further insight into pathogen-induced perturbations of indole GLS metabolism, we measured GLS in *cyp81F2-2* knock-out mutants that show loss of penetration resistance to *B. graminis* and enhanced susceptibility to *G. orontii*. 4MOI3M levels were significantly lower compared to wild-type independent of the treatment, confirming that CYP81F2 is the major CYP81F responsible for pathogen-induced 4MOI3M accumulation as well as for the establishment of basal 4MOI3M levels in whole leaves (Fig. 4). Total indole GLS levels were not significantly different in *cyp81F2-2* knock-out mutant and wild-type plants except for the *G. orontii* samples at 48 hpi, in which the knock-out mutant displayed higher levels than wild-type. Consequently, the decrease of total indole GLS levels observed in wild-type plants inoculated with *G. orontii* between 24 and 48 hpi was not detectable in the mutant. As expected for *cyp81F2-2*, I3M levels were overall higher than wild-type, but did not change dramatically in response to the treatment. This indicates that I3M is directly converted to 4MOI3M in the wild-type and that a threshold I3M concentration might exist that leads to feedback inhibition of indole GLS synthesis. To sum up, incompatible interactions with the non-adapted powdery mildew *B. graminis* appear to timely induce *de novo* I3M synthesis followed by CYP81F2-dependent production of 4MOI3M. In marked contrast, compatible interactions with the virulent pathogen *G. orontii* appear to correlate with reduced 4MOI3M synthesis, which may explain compatibility.

### Pathogen-induced epidermal SUR1 accumulation is absent in response to *G. orontii* infection

As 4MOI3M accumulation is absent at 24 hpi and 48 hpi with *G. orontii* despite prominent induction of *CYP83B1* and *CYP81F2* gene expression in the attacked epidermal cells as compared to those attacked by *B. graminis*, we set out to identify genes involved in indole GLS biosynthesis that might be perturbed in response to *G. orontii*. Among the biosynthetic enzymes, the C-S lyase SUR1 catalyzing the conversion of *S*-alkyl-thiohydroximates to thiohydroximates represented the most promising candidate for two reasons: First, the SUR1-catalyzed reaction is the only step in indole GLS synthesis lacking functional redundancy (Sønderby *et al*., 2010). Second, loss-of-function mutations of *SUR1* results in “pathway abortion” that manifests itself in the irreversible and spontaneous, intramolecular cyclization of the SUR1 substrate *S*-alkyl-thiohydroximate (Geu-Flores et al., 2009, 2011; Mikkelsen, Naur, & Halkier, 2004). To elucidate the dynamics of *SUR1* expression and localization in response to *G. orontii*, we inoculated three independent lines expressing *SUR1-mVenus* under the control of its endogenous promoter sequence and subsequently analyzed SUR1 accumulation by CLSM (Fig. 5). As an additional control, we inoculated the same set of fluorophore lines with *B. graminis* in parallel. SUR1-mVenus fluorescence was below the detection limit in the adaxial epidermis of unchallenged plants (Fig. 5a, while showing prominent fluorescence in vascular parenchyma cells as revealed by CLSM using cleared leaves (Fig. S1) and transverse sections of petioles (data not shown). Following inoculation with *G. orontii*, no SUR1-mVenus fluorescence was observed in epidermal cells at 24 hpi while weak fluorescence was detected at 48 hpi. By contrast, weak SUR1-mVenus fluorescence was observed at both 24 hpi and 48 hpi with *B. graminis*. Quantification of mean fluorescence intensities in attacked cells showed no significant differences for SUR1-mVenus accumulation in response to *G. orontii* (Fig. 5b. By contrast, SUR1-mVenus signal was significantly increased at 48 hpi with *B. graminis*. Hence, pathogen-induced SUR1 accumulation is absent in response to *G. orontii* infection. This finding is in line with the observed absence of 4MOI3M accumulation despite pathogen-induced accumulation of CYP83B1 and CYP81F2, and supports the idea that the indole core structure synthesis pathway is aborted.

**Figure 5:**
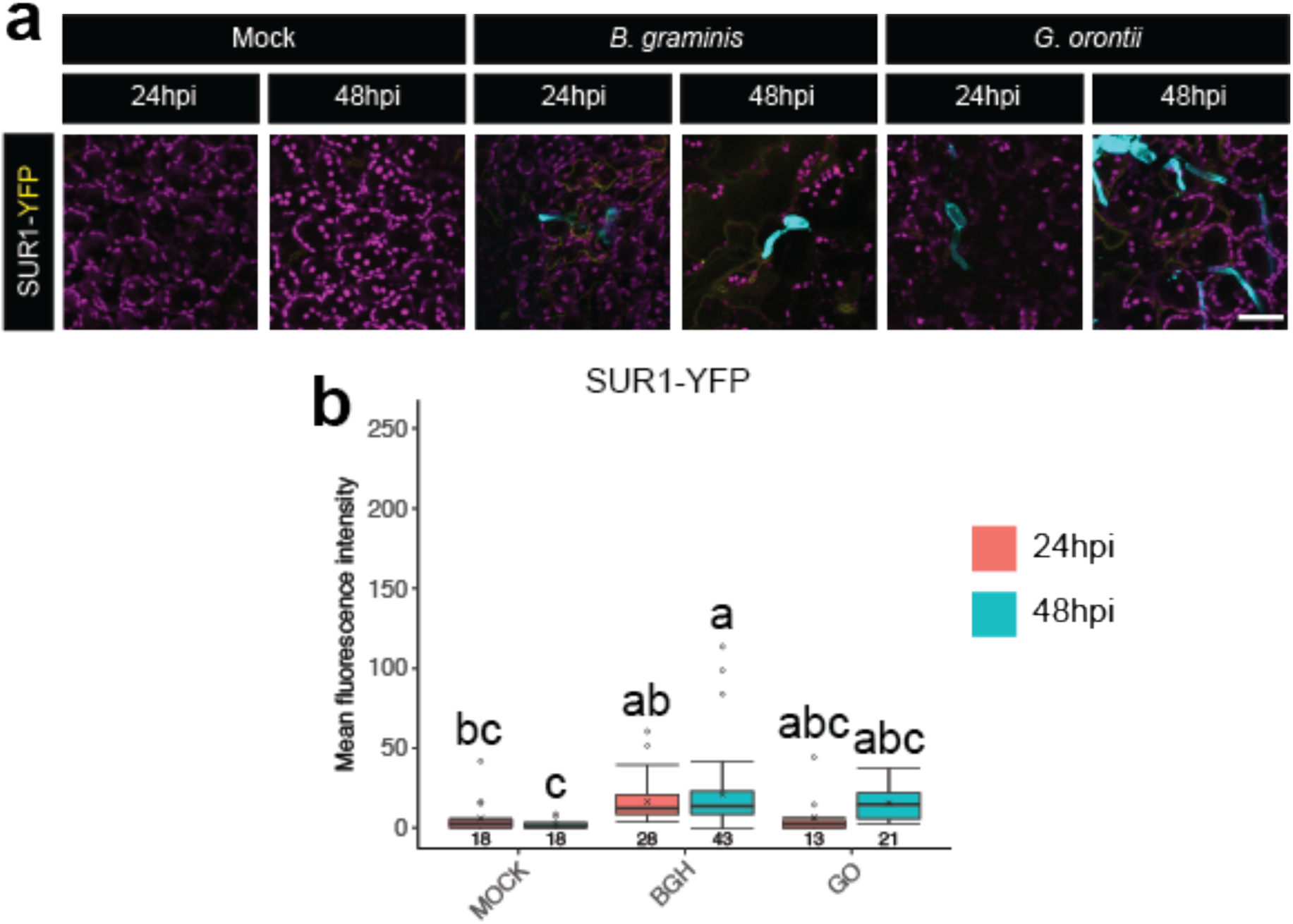
SUR1-YFP accumulation is induced by *B. graminis*, but not by *G. orontii*. (a) Representative confocal micrographs of plants expressing SUR1-YFP under the control of their native promoters at 24 hours post inoculation (hpi) and 48 hpi with *B. graminis* or *G. orontii* and following mock-treatment. All images represent z-projections through the adaxial epidermal cell layer. All images are overlays of YFP channel depicted in yellow, chlorophyll autofluorescence in magenta and calcofluor white staining in cyan. Scale bar = 50 µm. (b) Quantification of mean fluorescence intensities of SUR1-YFP following mock-treatment (MOCK) or inoculation with *B. graminis* (BGH) or *G. orontii* (GO) for 24 hpi (red boxes) and 48 hpi (blue boxes). Fluorescence was corrected by subtracting autofluorescence as determined in Col-0 plants (see Fig. S2). Results from three independent lines and three independent experiments were pooled. Individual boxplots show median (center line), mean (cross), first quartile (lower hinge), third quartile (upper hinge), whiskers (extending 1.5 times the inter-quartile range) and possible outliers (circles). Values below each box indicate the number of observations. Letters indicate significant differences between treatment × time point interactions as determined by two-way ANOVA (p<0.05) with Tukey HSD post-hoc test.

## Discussion

### The potential role of *PEN2* in post-invasive growth of adapted powdery mildew

Indole GLS-producing plants control numerous non-adapted and adapted plant pathogens via highly coordinated PEN2-mediated generation of toxic 4MOI3M hydrolysis products. Both pathogens used in this study have previously been shown to induce 4MOI3M accumulation, indicating increased flux from I3M to the PEN2 substrate 4MOI3M (Pawel Bednarek et al., 2009; Schön et al., 2013). Accordingly, expression of *CYP81F2* – the key enzyme for 4MOI3M synthesis – has been shown to be induced by both pathogens (Fuchs *et al*., 2016; Schön *et al*., 2013). However, *pen2-1* mutants were reported to be only impaired in non-host penetration resistance e.g. to incompatible interaction with *B. graminis*, as entry of virulent *G. orontii* was not affected at 24 hpi (Lipka *et al*., 2005). Nevertheless, we found that *G. orontii* displays significantly increased sporulation on *pen2-1* that was indistinguishable from that on *eds1-2* (Fig. 2a). This finding indicates that PEN2 is required to restrict post-invasive growth of *G. orontii*, which might be explained by a metabolic cost to detoxify products of PEN2 hydrolysis. Alternatively, PEN2 hydrolysis products might also react with endogenous plant compounds (e.g. amino acids) and thereby hamper the assimilatory process of the intruder. Furthermore, our results confirm that *pen2-1* mutants display enhanced *B. graminis* entry rates (Fig. 2b) (Lipka *et al*., 2005).

### Cell-autonomous induction of GLS biosynthesis upon attack

This study investigated the origin of I3M – the 4MOI3M precursor and substrate of the CYP81F2/PEN2 pathway – during the interaction of Arabidopsis with the powdery mildews *B. graminis* and *G. orontii*. We were able to detect 4MOI3M accumulation at both 24 hpi and 48 hpi with *B. graminis* (Fig. 4), despite the insignificance of CYP81F2 induction at 48 hpi (Fig. 3), indicating that the observed trend of CYP81F2 induction at 48 hpi is sufficient for 4MOI3M accumulation. Our results suggest that I3M is *de novo* synthesized as revealed by induction of CYP83B1 and SUR1 accumulation in the epidermis upon attack by *B. graminis* (Fig. 3 and Fig. 5). Similar results were obtained following inoculation with the biotrophic oomycete *Hyaloperonospora arabidopsidis* (Fig. S6). Induction of core structure synthesis is supported by the previous finding that γ-glutamylcysteine synthase, which is the enzyme catalyzing the first committed step in the biosynthesis of glutathione that is the sulfur donor in indole GLS synthesis, is not required for constitutive GLS accumulation, but for induction of indole GLS upon herbivory feeding and fungal attack (Schlaeppi, Abou-Mansour, Buchala, & Mauch, 2010; Schlaeppi, Bodenhausen, Buchala, Mauch, & Reymond, 2008).

### Transport of GLS during powdery mildew infection

In addition to the proposed cell-autonomous model for defence of epidermal cells against powdery mildews, multicellular defence systems might (co-)exist. Based on the vascular localization of GLS biosynthetic enzymes under normal growth conditions, one of our initial hypotheses was that core structure synthesis is induced in the vasculature upon attack and that I3M is subsequently transported to the attacked epidermal cells. This scenario would require signal transmission from epidermal cells to the sites of synthesis in the vasculature. Thereafter, GLS transport from vascular parenchyma cells in the phloem towards attacked epidermal cells might be facilitated by GTRs (Madsen *et al*., 2014). Alternatively, GLS might follow the symplasmic route via plasmodesmata (Ganusova & Burch-Smith, 2019) or be delivered via a combination of transmembrane and symplasmic transport.

We hypothesized that the characterized GLS importers GTR1-3 could be involved in import into attacked epidermal cells and/or export from distant organs. In the former case, GTRs would have to be localized to epidermal cells challenged by powdery mildews. Microscopy revealed that accumulation of neither GTR1 nor GTR2 is significantly induced in attacked cells, indicating that these transporters are not involved in defence against powdery mildews (Fig. 1). As the *gtr1gtr2* and *gtr1gtr2gtr3* mutants showed the same phenotype as wild-type in penetration resistance and sporulation of *B. graminis* and *G. orontii*, respectively (Fig. 2), we can exclude that the transporters have effects during important stages of disease development (i.e. time points other than 24 hpi and 48 hpi) not tested by microscopy. Moreover, comparing *gtr1gtr2* and *gtr1gtr2gtr3* mutants allowed to exclude an additive role of GTR3. Furthermore, interorgan redistribution of GLS upon attack such as e.g. enhanced root-to-shoot GLS translocation via downregulation or inactivation of GTRs in the root can be excluded. Although GTR-mediated GLS transport is not relevant for defence against powdery mildews, it is possible that other unknown transport proteins are involved. Furthermore, we did not address the role of plasmodesmata-mediated symplasmic transport of GLS towards attacked cells. One benefit of symplasmic GLS transport might be that it is regulated by sink strength, while drawbacks include that open plasmodesmata also allow intercellular transport of nutrients, effectors and fungal toxins. However, a previous study demonstrated that recognition of the fungal elicitor chitin limits the molecular flux through plasmodesmata (Faulkner *et al*., 2013). It is also noteworthy that powdery mildews exert pressure of 2-4 MPa during penetration (Micali, Göllner, Humphry, Consonni, & Panstruga, 2008; Tucker & Talbot, 2001). Independent of chitin perception, this pressure might be sufficient to trigger closure of plasmodesmata as demonstrated using pressure probes (Oparka & Prior, 1992).

### Remobilization of preformed GLS upon attack

Yet another hypothesis that would not require *de novo* GLS synthesis relies on remobilization of preformed I3M. GLS accumulate in vacuoles assuming that findings obtained by immunolocalization of GLS in *Brassica napus* can be translated to Arabidopsis (Kelly, Bones, & Rossiter, 1998). In rosette leaves, GLS concentrations are highest in the epidermis, especially at the leaf margins, and in S-cell that are located at the phloem cap. Cell-autonomous release of preformed and intracellularly stored I3M would thus require a mechanism for export from the vacuole, while remobilization from distant storage sites such as the S-cell would additionally employ pathways such as GTR-mediated transmembrane or plasmodesmata-mediated symplasmic transport. Arguing against remobilization, we observed that I3M pools were not depleted upon attempted penetration, indicating that I3M is compartmentalized and not accessible to ER-anchored CYP81F2 and mitochondria-anchored PEN2 (Fig. 4). However, the data presented in this study cannot absolutely exclude a GLS remobilization. Identification of a putative vacuolar GLS exporter will enable to directly address whether remobilization plays a role in these interactions.

### The role of indole GLS metabolism during PAMP-triggered immunity

Our results demonstrate that accumulation of CYP83B1 and CYP81F2 is induced upon powdery mildew infection (Fig. 3). Analysis of indole GLS biosynthesis gene expression revealed that *CYP81F2* is also highly induced by chitin, while *CYP83B1* and other genes encoding for enzymes in the indole GLS core structure pathway are not (Fig. S4). In addition, expression of the methyltransferases *IGMT1* and *IGMT2*, which catalyze 4-methoxylation of 4OHI3M, are highly induced upon chitin treatment (Fig. S4), indicating that side chain modification, but not core structure synthesis of I3M are elicited by chitin. In line with this finding, other studies suggested that only 4-substitution, but not *de novo* I3M synthesis is induced during plant-pathogen interactions (Pawel Bednarek et al., 2009; Iven et al., 2012). Our results using fluorophore-tagged proteins and live-cell microscopy contradict these earlier findings (Fig. 3 and Fig. 5). The results obtained in this study might be explained methodologically. Here, we utilized fluorophore-tagged protein fusions and thus collected data reflecting the protein level as compared to earlier studies that were investigating transcript abundance. It is feasible that enzymes involved in *de novo* I3M synthesis are regulated at the post-transcriptional level, while those required for side chain modifications are regulated transcriptionally. Furthermore, these earlier approaches might have been limited by the detection limit. Our results show that accumulation of core structure synthesis enzymes is not induced as high as those required for side chain modifications. Moreover, as only a subgroup of epidermal cells is attacked, the measurement of transcripts in whole leaves might not be able to resolve the induction of core structure synthesis enzymes. Similarly, the indole GLS concentration is supposedly solely induced in attacked epidermal cells. Hence, analysis of GLS or transcripts in whole leaves might be extremely diluted. We tried to overcome this issue by saturating the response using high numbers of conidiospores for inoculations. However, although the changes in GLS levels were significant, only slight differences were detected (Fig. 4). In regard to the massive induction of CYP81F2 (Fig. 3), the slight but significant changes of GLS indicate that the response is highly diluted, demonstrating that live-cell microscopy is a superior method to detect cell-autonomous processes and that pathogen-induced events might be masked in most transcriptomics, metabolomics and proteomics studies. The transcriptome of laser-capture microdissected haustorial complexes formed by *G. orontii* in Arabidopsis epidermal cells has been reported, but was able to detect only slight induction of indole GLS synthetic enzymes at 5 dpi (Chandran, Inada, Hather, Kleindt, & Wildermuth, 2010). In this study, the authors concluded that indole GLS synthesis might be delayed during the compatible interactions with *G. orontii* and that this delay might determine the compatibility. This finding is consistent with our result that 4MOI3M accumulates at 24 hpi and 48 hpi with *B. graminis*, but not *G. orontii* (Fig. 4). It is possible that 4MOI3M accumulates at later stages of infection with *G. orontii*, which would be in accordance with the effect of PEN2 during post-, but not pre-invasive growth of *G. orontii* (Fig. 2).

### *SUR1* as a potential powdery mildew effector target

Despite induced accumulation of both CYP83B1 and CYP81F2 during the interaction with *B. graminis* and *G. orontii*, 4MOI3M accumulates solely in response to *B. graminis*, suggesting that the indole GLS biosynthetic pathway is targeted by effectors of *G. orontii* (Fig. 3 and Fig. 4). Using CLSM, we show that induction of SUR1-mVenus accumulation is absent at 24 hpi and 48 hpi with *G. orontii*, but not with *B. graminis* (Fig. 5), indicating that SUR1 accumulation is repressed by *G. orontii*. It is also possible that SUR1 accumulation is not entirely absent during infection, but present at later stages of infection and thus only delayed. *SUR1* represents an optimal effector target due to the reported abortion of the indole GLS synthesis pathway in its absence. In absence of SUR1, *S*-alkyl-thiohydroximate will spontaneously and irreversibly cyclize (Geu-Flores et al., 2009, 2011; Mikkelsen et al., 2004). Furthermore, *SUR1* has no homologues and loss of *SUR1* function displays the most severe phenotype of all GLS biosynthetic enzymes as it results in seedling lethality due to auxin overaccumulation (Boerjan *et al*., 1995; Mikkelsen *et al*., 2004). By targeting *SUR1* via an effector, *G. orontii* might thus additionally profit from auxin overaccumulation, which results in loosening of the cell wall that in turn facilitates penetration (Dünser & Kleine-Vehn, 2015). Interestingly, SUR1 is the only core structure pathway enzyme that represents a target for endogenous miRNA (Kong, Li, Zhang, Jin, & Li, 2015). Targeting *SUR1* might thus be a straight-forward mechanism to regulate indole GLS synthesis and potentially also auxin synthesis that might be more advantageous as compared to non-selective regulation of all enzymes in the pathway via transcription factors such as *MYB34*, *MYB51* and *MYB122 (Frerigmann & Gigolashvili, 2014)*. Future studies should be directed towards identification and characterization of the hypothesized *G. orontii* effector and the role of the GLS core structure biosynthesis pathway as a potential breeding target for plant resistance of *Brassicaceae* crops.

### Accession numbers

*CYP81F2* (AT5G57220); *CYP83A1* (AT4G13770); *CYP83B1/SUR2* (AT4G31500); *EDR1* (AT1G08720); *EDS1* (AT3G48090); *NPF2.9/GTR3* (AT1G18880); *NPF2.10/GTR1* (AT3G47960); *NPF2.11/GTR2* (AT5G62680); *PEN2* (AT2G44490); *SUR1* (AT2G20610)

## Supporting Information

**Figure S1:** Tagged glucosinolate biosynthetic enzymes and transporters localize to cells of the vasculature under normal growth conditions.

**Figure S2:** Quantification of mean fluorescence intensities of GTR1-YFP and GTR2-mOrange2 following mock-treatment or inoculation with *B. graminis* or *G. orontii*.

**Figure S3:** Autofluorescence controls.

**Figure S4:** Transcripts abundance of indole GLS core structure synthesis and side chain modification in response to chitin as determined by microarray analysis.

**Figure S5:** Accumulation of aliphatic glucosinolates in wild-type and *cyp81f2* knock-out plants upon mock-, *B. graminis* and *G. orontii* treatment.

**Figure S6:** Cell-autonomous induction of CYP83B1 and CYP81F2 in response to *Hyaloperonospora arabidopsidis* infection.

**Appendix S1:** ANOVA results

