## Supplemental Data for "*De novo* indol-3-ylmethyl glucosinolate biosynthesis, and not long-distance transport, contributes to defence of Arabidopsis against powdery mildew"

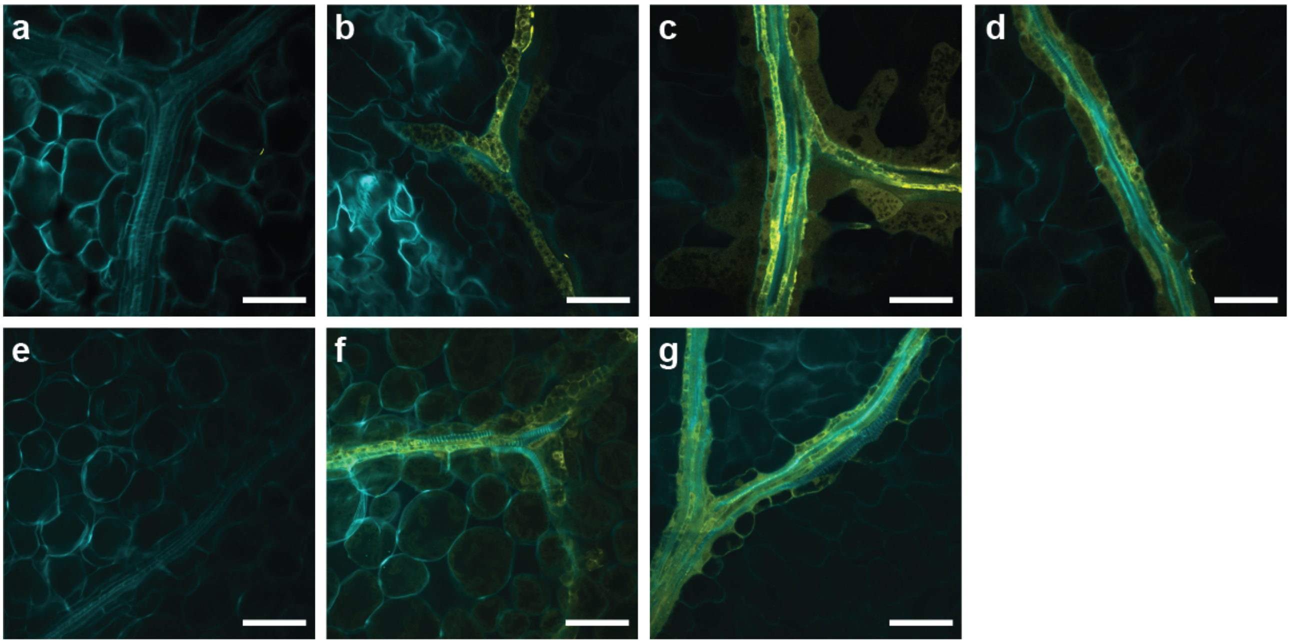


**Figure S1: Tagged glucosinolate biosynthetic enzymes and transporters localize to cells of the vasculature under normal growth conditions.** Confocal images of ClearSee-treated (a) Col-0 wild-type, (b) *CYP83A1-YFP*, (c) *CYP83B1-YFP*, (d) *SUR1-YFP*, (e) *gtr1gtr2*, (f) *GTR1-YFP/gtr1gtr2* and (g) *GTR2-mOrange2/gtr1gtr2* transgenic plants depicting fluorescence signal in yellow and calcofluor white staining of cell walls in cyan. (a-d) Images were acquired using YFP settings. (e-g) Images were acquired using mOrange settings. All images show vascular bundles of fifth leaves of 4-week-old plants. Photonmultiplier tubes were used for detection instead of HyD detectors. Scale bars = 50 µm.

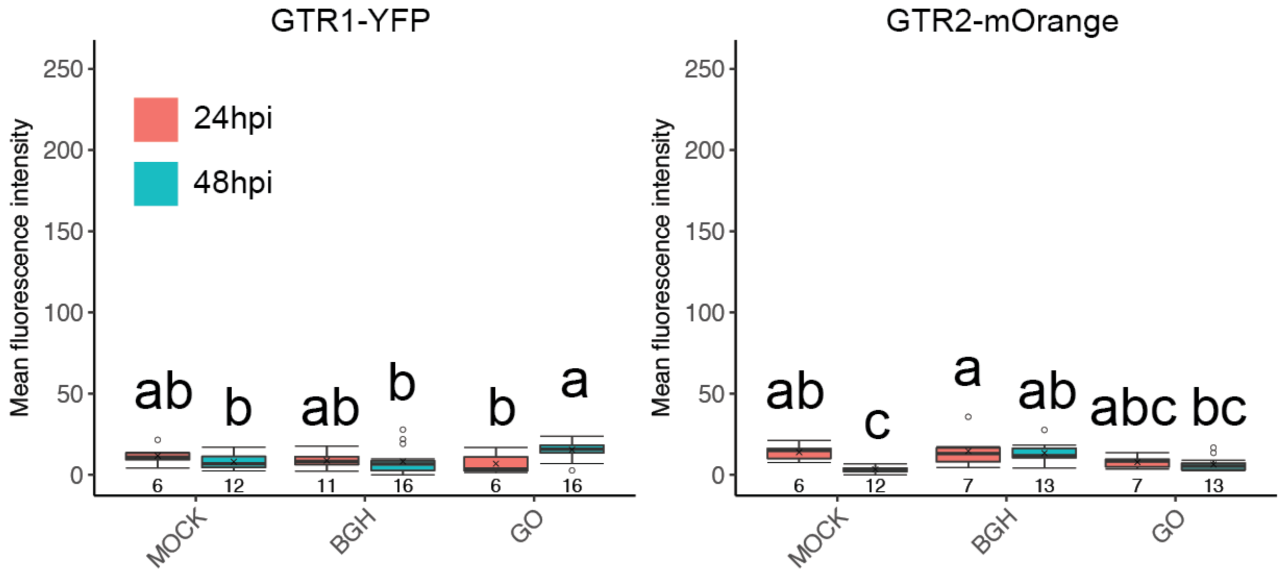


**Figure S2:** Quantification of mean fluorescence intensities of GTR1-YFP and GTR2-mOrange2 following mock-treatment (MOCK) or inoculation with *B. graminis* (BGH) or *G. orontii* (GO) for 24 hpi (red boxes) and 48 hpi (blue boxes). Fluorescence was corrected by subtracting autofluorescence as determined in Col-0 plants (see Fig. S3). Pooled data from two independent experiments are shown. Individual boxplots show median (center line), mean (cross), first quartile (lower hinge), third quartile (upper hinge), whiskers (extending 1.5 times the inter-quartile range) and possible outliers (circles). Values below each box indicate the number of observations. Letters indicate significant differences between treatment × timepoint interactions as determined by two-way ANOVA ($p<0.05$) with Tukey HSD post-hoc test.

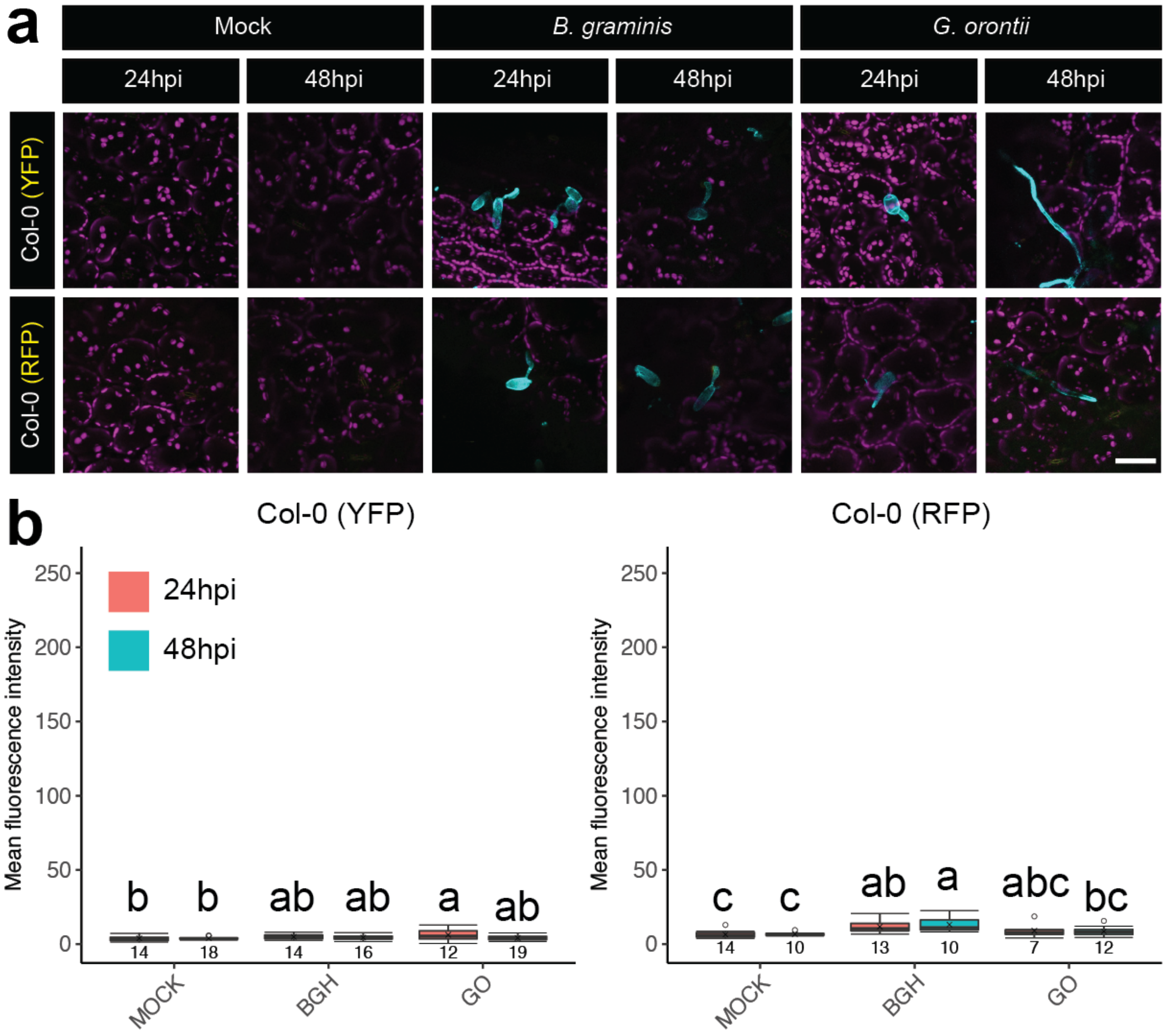


**Figure S3: Autofluorescence controls.** (a) Representative confocal micrographs of wild-type Col-0 plants imaged using YFP and RFP settings at 24 hours post inoculation (hpi) and 48 hpi with *B. graminis* or *G. orontii* and following mock-treatment. All images represent z-projections through the adaxial epidermal cell layer. All images are overlays of YFP or RFP channels both depicted in yellow, chlorophyll autofluorescence in magenta and calcofluor white staining in cyan. Scale bar = 50 µm. (b) Quantification of mean fluorescence intensities following mock-treatment (MOCK) or inoculation with *B. graminis* (BGH) or *G. orontii* (GO) for 24 hpi (red boxes) and 48 hpi (blue boxes). Pooled data from three independent experiments are shown. Individual boxplots show median (center line), mean (cross), first quartile (lower hinge), third quartile (upper hinge), whiskers (extending 1.5 times the inter-quartile range) and possible outliers (circles). Values below each box indicate the number of observations. Letters indicate significant differences between treatment × timepoint interactions as determined by two-way ANOVA ($p<0.05$) with Tukey HSD post-hoc test.

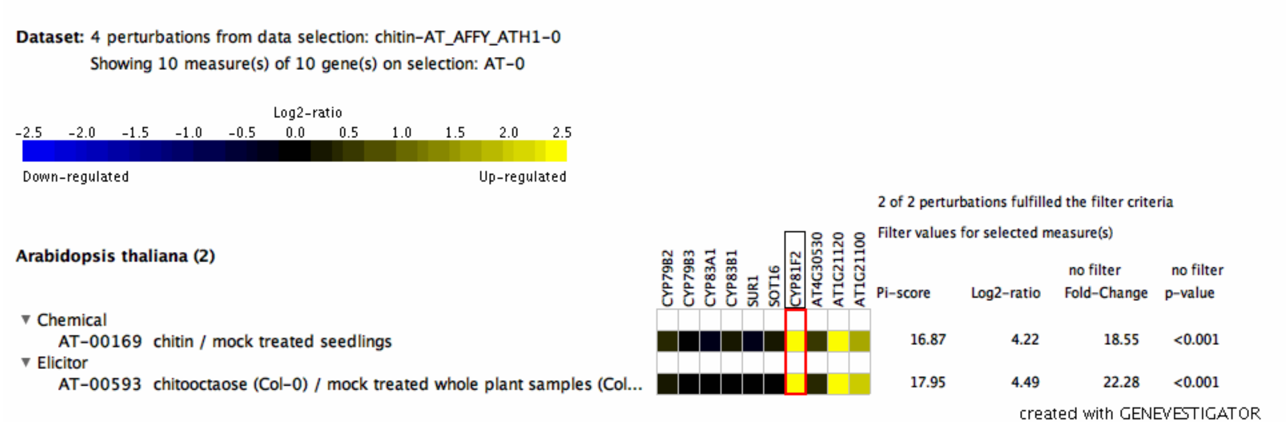


**Figure S4: Transcripts abundance of indole GLS core structure synthesis and side chain modification in response to chitin as determined by microarray analysis.**

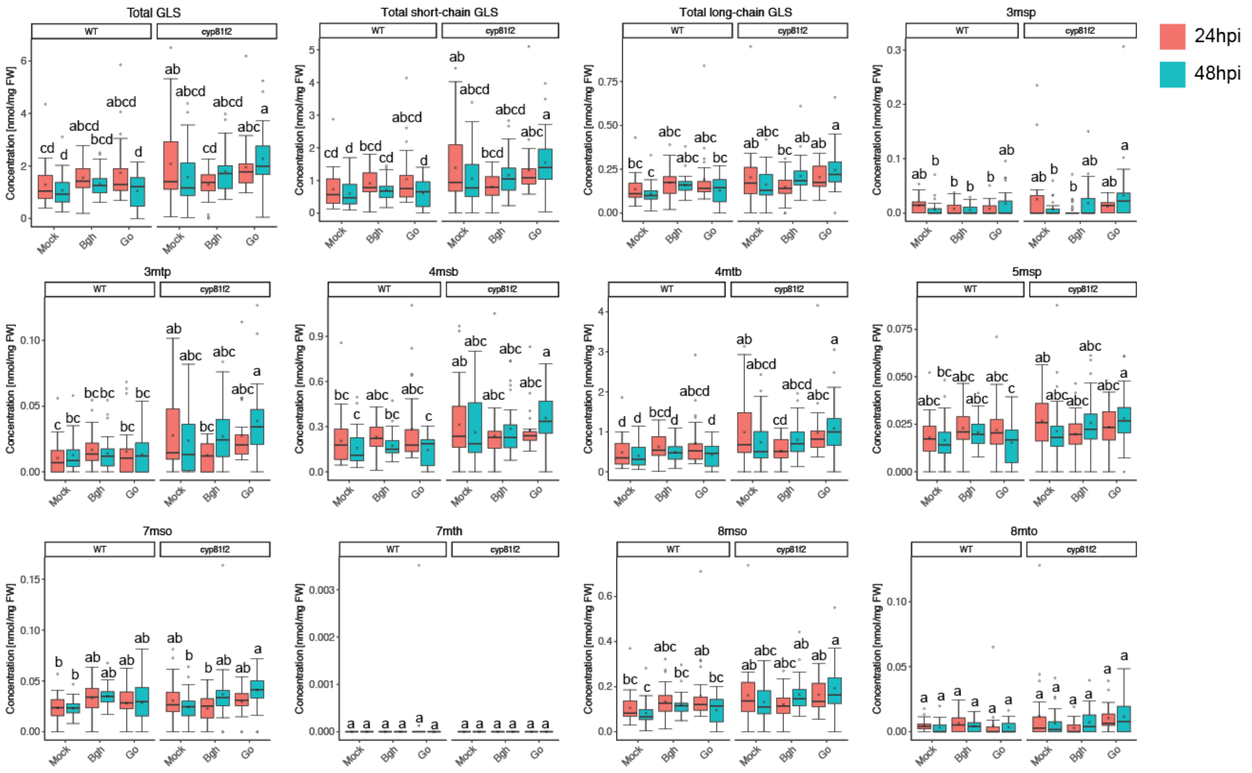


**Figure S5: Accumulation of aliphatic glucosinolates in wild-type and *cyp81f2* knock-out plants upon mock-, *B. graminis* and *G. orontii* treatment.** Quantification of total, total short-chain aliphatic (C3-C5), total long-chain aliphatic (C6-C8), 3msp, 3mtp, 4msb, 4mtb, 5msp, 7mso, 7mth, 8mso and 8mto glucosinolates in whole leaves of Col-0 (wild-type; WT) and *cyp81f2* mutant plants following mock-treatment (Mock) or inoculation with *B. graminis* (Bgh) or *G. orontii* (Go) for 24 hours (red boxes) and 48 hours (blue boxes). Pooled data from three independent experiments are shown. Individual boxplots show median (center line), mean (cross), first quartile (lower hinge), third quartile (upper hinge), whiskers (extending 1.5 times the inter-quartile range) and possible outliers (circles). Letters indicate significant differences between treatment × timepoint interactions as determined by two-way ANOVA ($p<0.05$; $n = 30$) with Tukey HSD post-hoc test.

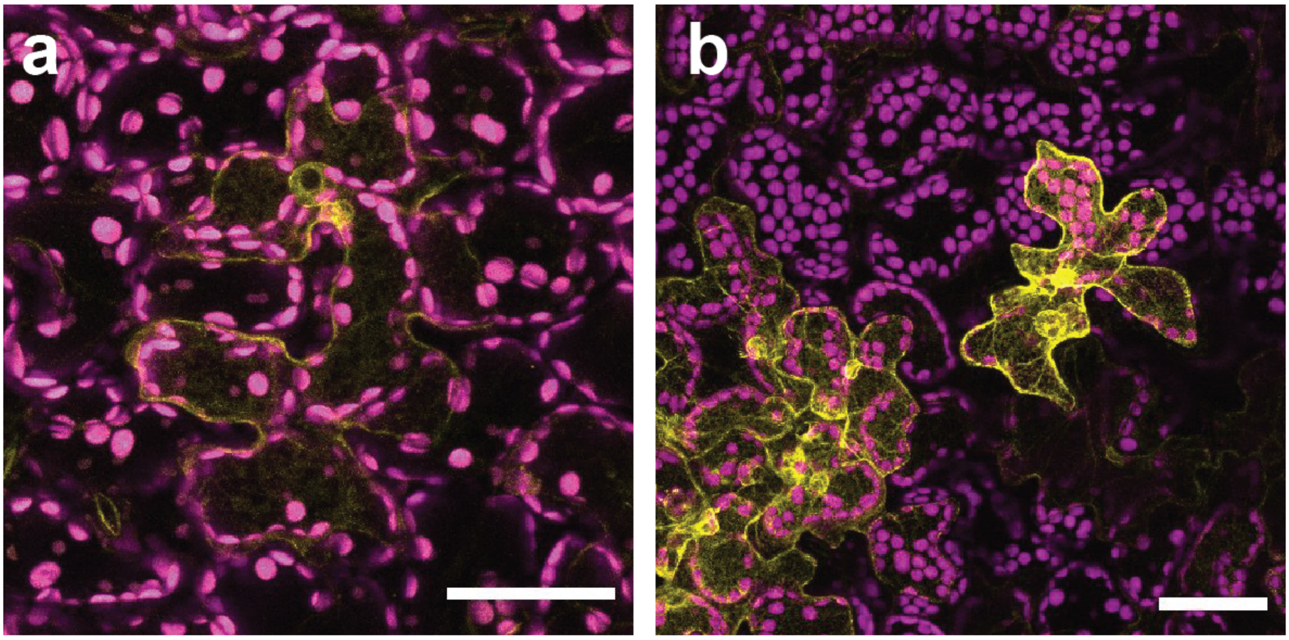


**Figure S6: Cell-autonomous induction of *CYP83B1* and *CYP81F2* in response to *Hyaloperonospora arabidopsidis* infection.** Representative maximum projections of z-stacks covering the epidermal cell layer of fifth leaves of (a) *CYP83B1-YFP* and (b) *CYP81F2-RFP* expressing transgenic plants at 96 hours post inoculated with *Hyaloperenospora arabidopsidis* NOCO2. YFP and RFP signals are shown in yellow. Chlorophyll autofluorescence is shown in magenta. Scale bars = 50 µm.

### Appendix S1: ANOVA results

#### Figure 2a:

| | Df | Sum Sq | Mean Sq | F value | Pr(>F) | |
|---|---|---|---|---|---|---|
| Genotype | 7 | 8.559e+13 | 1.223e+13 | 34.06 | < 2e-16 | *** |
| Experiment | 2 | 1.055e+13 | 5.275e+12 | 14.69 | 2.85e-06 | *** |
| Genotype:Experiment | 13 | 6.325e+13 | 4.866e+12 | 13.55 | 2.66e-16 | *** |
| Residuals | 93 | 3.339e+13 | 3.590e+11 | | | |

---
Signif. codes: 0 ‘***’ 0.001 ‘**’ 0.01 ‘*’ 0.05 ‘.’ 0.1 ‘ ’ 1

#### Figure 2b – Papillae:

| | Df | Sum Sq | Mean Sq | F value | Pr(>F) | |
|---|---|---|---|---|---|---|
| Genotype | 5 | 7536 | 1507.3 | 37.353 | 2.8e-13 | *** |
| Experiment | 2 | 413 | 206.7 | 5.123 | 0.01102 | * |
| Genotype:Experiment | 10 | 1194 | 119.4 | 2.960 | 0.00809 | ** |
| Residuals | 36 | 1453 | 40.4 | | | |

---
Signif. codes: 0 ‘***’ 0.001 ‘**’ 0.01 ‘*’ 0.05 ‘.’ 0.1 ‘ ’ 1

#### Figure 2b – Haustoria:

| | Df | Sum Sq | Mean Sq | F value | Pr(>F) | |
|---|---|---|---|---|---|---|
| Genotype | 5 | 2407.6 | 481.5 | 16.354 | 2.07e-08 | *** |
| Experiment | 2 | 343.6 | 171.8 | 5.835 | 0.00639 | ** |
| Genotype:Experiment | 10 | 882.6 | 88.3 | 2.998 | 0.00747 | ** |
| Residuals | 36 | 1060.0 | 29.4 | | | |

---
Signif. codes: 0 ‘***’ 0.001 ‘**’ 0.01 ‘*’ 0.05 ‘.’ 0.1 ‘ ’ 1

#### Figure 2b – Cell death:

| | Df | Sum Sq | Mean Sq | F value | Pr(>F) | |
|---|---|---|---|---|---|---|
| Genotype | 5 | 2092.6 | 418.5 | 46.695 | 9.53e-15 | *** |
| Experiment | 2 | 37.0 | 18.5 | 2.064 | 0.142 | |
| Genotype:Experiment | 10 | 86.6 | 8.7 | 0.966 | 0.489 | |
| Residuals | 36 | 322.7 | 9.0 | | | |

---
Signif. codes: 0 ‘***’ 0.001 ‘**’ 0.01 ‘*’ 0.05 ‘.’ 0.1 ‘ ’ 1

#### Figure 3b – CYP83A1-YFP:

| | Df | Sum Sq | Mean Sq | F value | Pr(>F) | |
|---|---|---|---|---|---|---|
| Experiment | 2 | 614.9 | 307.44 | 73.256 | < 2e-16 | *** |
| Genotype | 2 | 0.3 | 0.17 | 0.040 | 0.960644 | |
| Treatment | 2 | 38.4 | 19.22 | 4.579 | 0.011763 | * |
| Time | 1 | 53.3 | 53.25 | 12.689 | 0.000495 | *** |
| Experiment:Genotype | 2 | 23.2 | 11.60 | 2.763 | 0.066346 | . |
| Experiment:Treatment | 4 | 103.3 | 25.83 | 6.155 | 0.000131 | *** |
| Genotype:Treatment | 4 | 7.2 | 1.80 | 0.428 | 0.788402 | |
| Experiment:Time | 1 | 59.8 | 59.78 | 14.245 | 0.000232 | *** |
| Treatment:Time | 2 | 0.6 | 0.28 | 0.067 | 0.935406 | |
| Experiment:Genotype:Treatment | 3 | 11.1 | 3.69 | 0.880 | 0.452841 | |
| Experiment:Treatment:Time | 2 | 9.3 | 4.67 | 1.113 | 0.331451 | |
| Residuals | 148 | 621.1 | 4.20 | | | |

---
Signif. codes: 0 ‘***’ 0.001 ‘**’ 0.01 ‘*’ 0.05 ‘.’ 0.1 ‘ ’ 1

#### Figure 3b – CYP83B1-YFP:

| | Df | Sum Sq | Mean Sq | F value | Pr(>F) | |
|---|---|---|---|---|---|---|
| Experiment | 2 | 7825 | 3912 | 8.053 | 0.000419 | *** |
| Genotype | 2 | 40182 | 20091 | 41.353 | 4.96e-16 | *** |
| Treatment | 2 | 47546 | 23773 | 48.930 | < 2e-16 | *** |
| Time | 1 | 1571 | 1571 | 3.234 | 0.073447 | . |
| Experiment:Genotype | 2 | 13029 | 6515 | 13.409 | 3.14e-06 | *** |
| Experiment:Treatment | 4 | 3647 | 912 | 1.877 | 0.115436 | |
| Genotype:Treatment | 4 | 12862 | 3216 | 6.618 | 4.71e-05 | *** |
| Experiment:Time | 2 | 1486 | 743 | 1.529 | 0.219032 | |
| Genotype:Time | 1 | 187 | 187 | 0.385 | 0.535655 | |
| Treatment:Time | 2 | 1362 | 681 | 1.401 | 0.248413 | |
| Experiment:Genotype:Treatment | 4 | 5953 | 1488 | 3.063 | 0.017459 | * |
| Experiment:Genotype:Time | 1 | 807 | 807 | 1.661 | 0.198740 | |
| Experiment:Treatment:Time | 4 | 1517 | 379 | 0.781 | 0.538799 | |
| Genotype:Treatment:Time | 2 | 829 | 415 | 0.853 | 0.427329 | |
| Experiment:Genotype:Treatment:Time | 2 | 2008 | 1004 | 2.067 | 0.128980 | |
| Residuals | 226 | 109802 | 486 | | | |

---
Signif. codes: 0 '***' 0.001 '**' 0.01 '*' 0.05 '.' 0.1 ' ' 1

#### Figure 3b – CYP81F2-RFP:

| | Df | Sum Sq | Mean Sq | F value | Pr(>F) | |
|---|---|---|---|---|---|---|
| Experiment | 2 | 7061 | 3530 | 1.225 | 0.297726 | |
| Treatment | 2 | 118813 | 59406 | 20.621 | 2.66e-08 | *** |
| Time | 1 | 22305 | 22305 | 7.742 | 0.006378 | ** |
| Experiment:Treatment | 4 | 74755 | 18689 | 6.487 | 0.000103 | *** |
| Treatment:Time | 2 | 7106 | 3553 | 1.233 | 0.295437 | |
| Residuals | 107 | 308259 | 2881 | | | |

---
Signif. codes: 0 '***' 0.001 '**' 0.01 '*' 0.05 '.' 0.1 ' ' 1

#### Figure 4 – Total indole GLS:

| | Sum Sq | Df | F value | Pr(>F) | |
|---|---|---|---|---|---|
| Replicate | 1.0133 | 2 | 16.6037 | 1.412e-07 | *** |
| Genotype | 0.0000 | 1 | 0.0009 | 0.9766042 | |
| Treatment | 0.0391 | 2 | 0.6402 | 0.5278605 | |
| Time | 0.2352 | 1 | 7.7092 | 0.0058284 | ** |
| Replicate:Genotype | 0.2809 | 2 | 4.6027 | 0.0107196 | * |
| Replicate:Treatment | 0.2540 | 4 | 2.0814 | 0.0830971 | . |
| Genotype:Treatment | 0.2785 | 2 | 4.5629 | 0.0111415 | * |
| Replicate:Time | 0.2419 | 2 | 3.9638 | 0.0199615 | * |
| Genotype:Time | 0.1663 | 1 | 5.4488 | 0.0202206 | * |
| Treatment:Time | 0.3188 | 2 | 5.2243 | 0.0058684 | ** |
| Replicate:Genotype:Treatment | 0.1727 | 4 | 1.4152 | 0.2287221 | |
| Replicate:Genotype:Time | 0.0914 | 2 | 1.4973 | 0.2253477 | |
| Replicate:Treatment:Time | 0.1233 | 4 | 1.0103 | 0.4022390 | |
| Genotype:Treatment:Time | 0.4651 | 2 | 7.6216 | 0.0005872 | *** |
| Replicate:Genotype:Treatment:Time | 0.1047 | 4 | 0.8580 | 0.4895324 | |
| Residuals | 9.4591 | 310 | | | |

---
Signif. codes: 0 '***' 0.001 '**' 0.01 '*' 0.05 '.' 0.1 ' ' 1

#### Figure 4 – I3M:

| | Sum Sq | Df | F value | Pr(>F) | |
|---|---|---|---|---|---|
| Replicate | 0.6698 | 2 | 17.1212 | 8.853e-08 | *** |
| Genotype | 1.9411 | 1 | 99.2336 | < 2.2e-16 | *** |
| Treatment | 0.2686 | 2 | 6.8644 | 0.0012104 | ** |
| Time | 0.1374 | 1 | 7.0255 | 0.0084484 | ** |
| Replicate:Genotype | 0.1590 | 2 | 4.0638 | 0.0181066 | * |
| Replicate:Treatment | 0.0678 | 4 | 0.8663 | 0.4844161 | |
| Genotype:Treatment | 0.0379 | 2 | 0.9683 | 0.3808849 | |
| Replicate:Time | 0.1200 | 2 | 3.0670 | 0.0479778 | * |
| Genotype:Time | 0.1157 | 1 | 5.9161 | 0.0155685 | * |
| Treatment:Time | 0.1711 | 2 | 4.3727 | 0.0134052 | * |
| Replicate:Genotype:Treatment | 0.0641 | 4 | 0.8195 | 0.5134933 | |
| Replicate:Genotype:Time | 0.1171 | 2 | 2.9935 | 0.0515609 | . |
| Replicate:Treatment:Time | 0.1714 | 4 | 2.1906 | 0.0699353 | . |
| Genotype:Treatment:Time | 0.3298 | 2 | 8.4306 | 0.0002721 | *** |
| Replicate:Genotype:Treatment:Time | 0.0412 | 4 | 0.5267 | 0.7162195 | |
| Residuals | 6.0639 | 310 | | | |

---
Signif. codes: 0 '***' 0.001 '**' 0.01 '*' 0.05 '.' 0.1 ' ' 1

#### Figure 4 – 1MOI3M:

| | Sum Sq | Df | F value | Pr(>F) | |
|---|---|---|---|---|---|
| Replicate | 0.008571 | 2 | 5.6518 | 0.003883 | ** |
| Genotype | 0.000004 | 1 | 0.0056 | 0.940488 | |
| Treatment | 0.005512 | 2 | 3.6348 | 0.027521 | * |
| Time | 0.001255 | 1 | 1.6556 | 0.199153 | |
| Replicate:Genotype | 0.002870 | 2 | 1.8926 | 0.152419 | |
| Replicate:Treatment | 0.003466 | 4 | 1.1428 | 0.336378 | |
| Genotype:Treatment | 0.001129 | 2 | 0.7441 | 0.475994 | |
| Replicate:Time | 0.001383 | 2 | 0.9122 | 0.402722 | |
| Genotype:Time | 0.000013 | 1 | 0.0168 | 0.896814 | |
| Treatment:Time | 0.001021 | 2 | 0.6734 | 0.510736 | |
| Replicate:Genotype:Treatment | 0.002492 | 4 | 0.8215 | 0.512237 | |
| Replicate:Genotype:Time | 0.000319 | 2 | 0.2101 | 0.810581 | |
| Replicate:Treatment:Time | 0.003306 | 4 | 1.0900 | 0.361497 | |
| Genotype:Treatment:Time | 0.001046 | 2 | 0.6900 | 0.502332 | |
| Replicate:Genotype:Treatment:Time | 0.001808 | 4 | 0.5962 | 0.665640 | |
| Residuals | 0.235064 | 310 | | | |

---
Signif. codes: 0 '***' 0.001 '**' 0.01 '*' 0.05 '.' 0.1 ' ' 1

#### Figure 4 – 4MOI3M:

| | Sum Sq | Df | F value | Pr(>F) | |
|---|---|---|---|---|---|
| Replicate | 0.00915 | 2 | 1.8986 | 0.151514 | |
| Genotype | 1.92114 | 1 | 797.1909 | < 2.2e-16 | *** |
| Treatment | 0.20762 | 2 | 43.0765 | < 2.2e-16 | *** |
| Time | 0.00622 | 1 | 2.5809 | 0.109180 | |
| Replicate:Genotype | 0.03347 | 2 | 6.9451 | 0.001120 | ** |
| Replicate:Treatment | 0.08102 | 4 | 8.4050 | 1.894e-06 | *** |
| Genotype:Treatment | 0.29297 | 2 | 60.7852 | < 2.2e-16 | *** |
| Replicate:Time | 0.01188 | 2 | 2.4640 | 0.086760 | . |
| Genotype:Time | 0.00506 | 1 | 2.1000 | 0.148315 | |
| Treatment:Time | 0.01559 | 2 | 3.2353 | 0.040681 | * |
| Replicate:Genotype:Treatment | 0.09739 | 4 | 10.1030 | 1.065e-07 | *** |
| Replicate:Genotype:Time | 0.00354 | 2 | 0.7349 | 0.480369 | |
| Replicate:Treatment:Time | 0.03459 | 4 | 3.5888 | 0.007051 | ** |
| Genotype:Treatment:Time | 0.01198 | 2 | 2.4856 | 0.084930 | . |
| Replicate:Genotype:Treatment:Time | 0.02142 | 4 | 2.2218 | 0.066557 | . |
| Residuals | 0.74707 | 310 | | | |

---
Signif. codes: 0 '***' 0.001 '**' 0.01 '*' 0.05 '.' 0.1 ' ' 1

#### Figure 5b – SUR1-YFP:

| | Df | Sum Sq | Mean Sq | F value | Pr(>F) | |
|---|---|---|---|---|---|---|
| Experiment | 2 | 7323 | 3661 | 24.414 | 1.34e-09 | *** |
| Genotype | 1 | 142 | 142 | 0.946 | 0.3328 | |
| Treatment | 2 | 7169 | 3585 | 23.903 | 1.93e-09 | *** |
| Time | 1 | 479 | 479 | 3.196 | 0.0764 | . |
| Experiment:Genotype | 1 | 708 | 708 | 4.723 | 0.0318 | * |
| Experiment:Treatment | 4 | 5293 | 1323 | 8.823 | 2.88e-06 | *** |
| Genotype:Treatment | 2 | 839 | 419 | 2.796 | 0.0651 | . |
| Experiment:Time | 1 | 104 | 104 | 0.696 | 0.4060 | |
| Genotype:Time | 1 | 33 | 33 | 0.220 | 0.6398 | |
| Treatment:Time | 2 | 1016 | 508 | 3.387 | 0.0371 | * |
| Experiment:Genotype:Treatment | 2 | 388 | 194 | 1.292 | 0.2785 | |
| Experiment:Treatment:Time | 1 | 4 | 4 | 0.028 | 0.8673 | |
| Genotype:Treatment:Time | 2 | 104 | 52 | 0.348 | 0.7071 | |
| Residuals | 118 | 17696 | 150 | | | |

---
Signif. codes: 0 '***' 0.001 '**' 0.01 '*' 0.05 '.' 0.1 ' ' 1

#### Figure S2 – GTR1-YFP:

| | Df | Sum Sq | Mean Sq | F value | Pr(>F) | |
|---|---|---|---|---|---|---|
| Experiment | 1 | 520.9 | 520.9 | 18.846 | 5.75e-05 | *** |
| Treatment | 2 | 228.3 | 114.2 | 4.130 | 0.02104 | * |
| Time | 1 | 66.1 | 66.1 | 2.392 | 0.12736 | |
| Experiment:Treatment | 2 | 124.7 | 62.4 | 2.256 | 0.11386 | |
| Treatment:Time | 2 | 295.9 | 147.9 | 5.352 | 0.00736 | ** |
| Residuals | 58 | 1603.2 | 27.6 | | | |

---
Signif. codes: 0 '***' 0.001 '**' 0.01 '*' 0.05 '.' 0.1 ' ' 1

#### Figure S2 – GTR2-mOrange2:

| | Df | Sum Sq | Mean Sq | F value | Pr(>F) | |
|---|---|---|---|---|---|---|
| Experiment | 1 | 8.9 | 8.9 | 0.290 | 0.592365 | |
| Treatment | 2 | 646.4 | 323.2 | 10.544 | 0.000155 | *** |
| Time | 1 | 236.2 | 236.2 | 7.706 | 0.007773 | ** |
| Experiment:Treatment | 2 | 178.1 | 89.1 | 2.905 | 0.064204 | . |
| Treatment:Time | 2 | 107.3 | 53.7 | 1.750 | 0.184388 | |
| Residuals | 49 | 1502.1 | 30.7 | | | |

---
Signif. codes: 0 '***' 0.001 '**' 0.01 '*' 0.05 '.' 0.1 ' ' 1

#### Figure S3b – Col-0 (YFP):

| | Df | Sum Sq | Mean Sq | F value | Pr(>F) | |
|---|---|---|---|---|---|---|
| Experiment | 2 | 44.98 | 22.488 | 6.630 | 0.00214 | ** |
| Treatment | 2 | 15.47 | 7.737 | 2.281 | 0.10862 | |
| Time | 1 | 18.69 | 18.692 | 5.511 | 0.02131 | * |
| Experiment:Treatment | 4 | 39.76 | 9.940 | 2.931 | 0.02560 | * |
| Treatment:Time | 1 | 0.08 | 0.085 | 0.025 | 0.87463 | |
| Residuals | 82 | 278.13 | 3.392 | | | |

---
Signif. codes: 0 '***' 0.001 '**' 0.01 '*' 0.05 '.' 0.1 ' ' 1

#### Figure S3b – Col-0 (RFP):

| | Df | Sum Sq | Mean Sq | F value | Pr(>F) | |
|---|---|---|---|---|---|---|
| Experiment | 2 | 3.3 | 1.63 | 0.130 | 0.878 | |
| Treatment | 2 | 403.5 | 201.76 | 16.175 | 2.73e-06 | *** |
| Experiment:Treatment | 4 | 71.5 | 17.88 | 1.433 | 0.235 | |
| Residuals | 57 | 711.0 | 12.47 | | | |

---
Signif. codes: 0 '***' 0.001 '**' 0.01 '*' 0.05 '.' 0.1 ' ' 1

#### Figure S5 – Total GLS:

| | Sum Sq | Df | F value | Pr(>F) | |
|---|---|---|---|---|---|
| Replicate | 56.812 | 2 | 45.7897 | < 2.2e-16 | *** |
| Genotype | 22.385 | 1 | 36.0840 | 5.291e-09 | *** |
| Treatment | 6.027 | 2 | 4.8581 | 0.008367 | ** |
| Time | 3.194 | 1 | 5.1487 | 0.023951 | * |
| Replicate:Genotype | 1.292 | 2 | 1.0416 | 0.354139 | |
| Replicate:Treatment | 9.508 | 4 | 3.8316 | 0.004686 | ** |
| Genotype:Treatment | 7.462 | 2 | 6.0143 | 0.002738 | ** |
| Replicate:Time | 3.370 | 2 | 2.7158 | 0.067725 | . |
| Genotype:Time | 3.527 | 1 | 5.6847 | 0.017714 | * |
| Treatment:Time | 4.903 | 2 | 3.9518 | 0.020196 | * |
| Replicate:Genotype:Treatment | 4.457 | 4 | 1.7960 | 0.129441 | |
| Replicate:Genotype:Time | 4.909 | 2 | 3.9566 | 0.020103 | * |
| Replicate:Treatment:Time | 7.147 | 4 | 2.8803 | 0.022914 | * |
| Genotype:Treatment:Time | 6.149 | 2 | 4.9560 | 0.007609 | ** |
| Replicate:Genotype:Treatment:Time | 1.906 | 4 | 0.7679 | 0.546759 | |
| Residuals | 192.310 | 310 | | | |

---
Signif. codes: 0 '***' 0.001 '**' 0.01 '*' 0.05 '.' 0.1 ' ' 1

#### Figure S5 – Total short-chain GLS:

| | Sum Sq | Df | F value | Pr(>F) | |
|---|---|---|---|---|---|
| Replicate | 37.977 | 2 | 58.1242 | < 2.2e-16 | *** |
| Genotype | 18.321 | 1 | 56.0813 | 7.285e-13 | *** |
| Treatment | 4.159 | 2 | 6.3654 | 0.001953 | ** |
| Time | 1.453 | 1 | 4.4490 | 0.035723 | * |
| Replicate:Genotype | 1.747 | 2 | 2.6738 | 0.070584 | . |
| Replicate:Treatment | 5.391 | 4 | 4.1252 | 0.002852 | ** |
| Genotype:Treatment | 3.895 | 2 | 5.9617 | 0.002880 | ** |
| Replicate:Time | 1.329 | 2 | 2.0348 | 0.132450 | |
| Genotype:Time | 1.517 | 1 | 4.6434 | 0.031943 | * |
| Treatment:Time | 2.023 | 2 | 3.0957 | 0.046643 | * |
| Replicate:Genotype:Treatment | 2.423 | 4 | 1.8544 | 0.118337 | |
| Replicate:Genotype:Time | 2.968 | 2 | 4.5430 | 0.011359 | * |
| Replicate:Treatment:Time | 4.782 | 4 | 3.6596 | 0.006260 | ** |
| Genotype:Treatment:Time | 2.653 | 2 | 4.0606 | 0.018165 | * |
| Replicate:Genotype:Treatment:Time | 1.219 | 4 | 0.9331 | 0.444944 | |
| Residuals | 101.274 | 310 | | | |

---
Signif. codes: 0 '***' 0.001 '**' 0.01 '*' 0.05 '.' 0.1 ' ' 1

#### Figure S5 – Total long-chain GLS:

| | Sum Sq | Df | F value | Pr(>F) | |
|---|---|---|---|---|---|
| Replicate | 0.24972 | 2 | 14.6564 | 8.280e-07 | *** |
| Genotype | 0.19882 | 1 | 23.3384 | 2.139e-06 | *** |
| Treatment | 0.09318 | 2 | 5.4688 | 0.0046329 | ** |
| Time | 0.00859 | 1 | 1.0077 | 0.3162305 | |
| Replicate:Genotype | 0.03117 | 2 | 1.8294 | 0.1622344 | |
| Replicate:Treatment | 0.19572 | 4 | 5.7434 | 0.0001801 | *** |
| Genotype:Treatment | 0.05368 | 2 | 3.1507 | 0.0441965 | * |
| Replicate:Time | 0.11037 | 2 | 6.4779 | 0.0017534 | ** |
| Genotype:Time | 0.05462 | 1 | 6.4111 | 0.0118346 | * |
| Treatment:Time | 0.05527 | 2 | 3.2440 | 0.0403367 | * |
| Replicate:Genotype:Treatment | 0.04442 | 4 | 1.3035 | 0.2686115 | |
| Replicate:Genotype:Time | 0.04516 | 2 | 2.6502 | 0.0722375 | . |
| Replicate:Treatment:Time | 0.04769 | 4 | 1.3995 | 0.2339963 | |
| Genotype:Treatment:Time | 0.05693 | 2 | 3.3410 | 0.0366806 | * |
| Replicate:Genotype:Treatment:Time | 0.02443 | 4 | 0.7171 | 0.5807996 | |
| Residuals | 2.64095 | 310 | | | |

---
Signif. codes: 0 ‘***’ 0.001 ‘**’ 0.01 ‘*’ 0.05 ‘.’ 0.1 ‘ ’ 1

#### Figure S5 – 3mtp:

| | Sum Sq | Df | F value | Pr(>F) | |
|---|---|---|---|---|---|
| Replicate | 0.024944 | 2 | 44.6440 | < 2.2e-16 | *** |
| Genotype | 0.013632 | 1 | 48.7946 | 1.74e-11 | *** |
| Treatment | 0.002665 | 2 | 4.7706 | 0.009108 | ** |
| Time | 0.000359 | 1 | 1.2839 | 0.258051 | |
| Replicate:Genotype | 0.002936 | 2 | 5.2538 | 0.005703 | ** |
| Replicate:Treatment | 0.001823 | 4 | 1.6318 | 0.166044 | |
| Genotype:Treatment | 0.003544 | 2 | 6.3424 | 0.001997 | ** |
| Replicate:Time | 0.001408 | 2 | 2.5191 | 0.082182 | . |
| Genotype:Time | 0.000758 | 1 | 2.7126 | 0.100571 | |
| Treatment:Time | 0.000708 | 2 | 1.2666 | 0.283233 | |
| Replicate:Genotype:Treatment | 0.002760 | 4 | 2.4701 | 0.044721 | * |
| Replicate:Genotype:Time | 0.001174 | 2 | 2.1013 | 0.124032 | |
| Replicate:Treatment:Time | 0.004122 | 4 | 3.6889 | 0.005959 | ** |
| Genotype:Treatment:Time | 0.001987 | 2 | 3.5571 | 0.029693 | * |
| Replicate:Genotype:Treatment:Time | 0.002006 | 4 | 1.7952 | 0.129603 | |
| Residuals | 0.086604 | 310 | | | |

---
Signif. codes: 0 ‘***’ 0.001 ‘**’ 0.01 ‘*’ 0.05 ‘.’ 0.1 ‘ ’ 1

#### Figure S5 – 3msp:

| | Sum Sq | Df | F value | Pr(>F) | |
|---|---|---|---|---|---|
| Replicate | 0.003261 | 2 | 2.2795 | 0.1040444 | |
| Genotype | 0.003847 | 1 | 5.3786 | 0.0210355 | * |
| Treatment | 0.003811 | 2 | 2.6644 | 0.0712365 | . |
| Time | 0.000128 | 1 | 0.1794 | 0.6722254 | |
| Replicate:Genotype | 0.003969 | 2 | 2.7745 | 0.0639315 | . |
| Replicate:Treatment | 0.004343 | 4 | 1.5180 | 0.1967346 | |
| Genotype:Treatment | 0.000272 | 2 | 0.1903 | 0.8267845 | |
| Replicate:Time | 0.010129 | 2 | 7.0811 | 0.0009838 | *** |
| Genotype:Time | 0.000252 | 1 | 0.3521 | 0.5533792 | |
| Treatment:Time | 0.010389 | 2 | 7.2626 | 0.0008271 | *** |
| Replicate:Genotype:Treatment | 0.005244 | 4 | 1.8329 | 0.1223085 | |
| Replicate:Genotype:Time | 0.003491 | 2 | 2.4408 | 0.0887653 | . |
| Replicate:Treatment:Time | 0.002749 | 4 | 0.9610 | 0.4291375 | |
| Genotype:Treatment:Time | 0.002866 | 2 | 2.0035 | 0.1366113 | |
| Replicate:Genotype:Treatment:Time | 0.002034 | 4 | 0.7111 | 0.5848526 | |
| Residuals | 0.221721 | 310 | | | |

---
Signif. codes: 0 ‘***’ 0.001 ‘**’ 0.01 ‘*’ 0.05 ‘.’ 0.1 ‘ ’ 1

#### Figure S5 – 4mtb:

| | Sum Sq | Df | F value | Pr(>F) | |
|---|---|---|---|---|---|
| Replicate | 18.726 | 2 | 51.3342 | < 2.2e-16 | *** |
| Genotype | 10.065 | 1 | 55.1838 | 1.073e-12 | *** |
| Treatment | 2.564 | 2 | 7.0284 | 0.0010346 | ** |
| Time | 0.764 | 1 | 4.1866 | 0.0415884 | * |
| Replicate:Genotype | 1.046 | 2 | 2.8661 | 0.0584285 | . |
| Replicate:Treatment | 3.198 | 4 | 4.3829 | 0.0018413 | ** |
| Genotype:Treatment | 2.715 | 2 | 7.4440 | 0.0006956 | *** |
| Replicate:Time | 0.519 | 2 | 1.4226 | 0.2426643 | |
| Genotype:Time | 0.533 | 1 | 2.9238 | 0.0882847 | . |
| Treatment:Time | 1.309 | 2 | 3.5879 | 0.0288110 | * |
| Replicate:Genotype:Treatment | 1.691 | 4 | 2.3182 | 0.0570732 | . |
| Replicate:Genotype:Time | 1.300 | 2 | 3.5650 | 0.0294635 | * |
| Replicate:Treatment:Time | 2.529 | 4 | 3.4664 | 0.0086570 | ** |
| Genotype:Treatment:Time | 1.376 | 2 | 3.7719 | 0.0240718 | * |
| Replicate:Genotype:Treatment:Time | 1.061 | 4 | 1.4546 | 0.2159494 | |
| Residuals | 56.542 | 310 | | | |

---
Signif. codes:  0 ‘***’ 0.001 ‘**’ 0.01 ‘*’ 0.05 ‘.’ 0.1 ‘ ’ 1

#### Figure S5 – 4msb:

| | Sum Sq | Df | F value | Pr(>F) | |
|---|---|---|---|---|---|
| Replicate | 2.6058 | 2 | 68.4862 | < 2.2e-16 | *** |
| Genotype | 0.7809 | 1 | 41.0460 | 5.516e-10 | *** |
| Treatment | 0.1076 | 2 | 2.8274 | 0.060693 | . |
| Time | 0.1248 | 1 | 6.5602 | 0.010902 | * |
| Replicate:Genotype | 0.0363 | 2 | 0.9532 | 0.386623 | |
| Replicate:Treatment | 0.2777 | 4 | 3.6487 | 0.006375 | ** |
| Genotype:Treatment | 0.0519 | 2 | 1.3648 | 0.256954 | |
| Replicate:Time | 0.1164 | 2 | 3.0582 | 0.048393 | * |
| Genotype:Time | 0.1882 | 1 | 9.8905 | 0.001822 | ** |
| Treatment:Time | 0.0475 | 2 | 1.2493 | 0.288140 | |
| Replicate:Genotype:Treatment | 0.1005 | 4 | 1.3201 | 0.262333 | |
| Replicate:Genotype:Time | 0.2223 | 2 | 5.8426 | 0.003230 | ** |
| Replicate:Treatment:Time | 0.3269 | 4 | 4.2962 | 0.002134 | ** |
| Genotype:Treatment:Time | 0.1557 | 2 | 4.0927 | 0.017605 | * |
| Replicate:Genotype:Treatment:Time | 0.0227 | 4 | 0.2986 | 0.878780 | |
| Residuals | 5.8975 | 310 | | | |

---
Signif. codes:  0 ‘***’ 0.001 ‘**’ 0.01 ‘*’ 0.05 ‘.’ 0.1 ‘ ’ 1

#### Figure S5 – 5msp:

| | Sum Sq | Df | F value | Pr(>F) | |
|---|---|---|---|---|---|
| Replicate | 0.002477 | 2 | 8.7348 | 0.0002040 | *** |
| Genotype | 0.002056 | 1 | 14.4980 | 0.0001691 | *** |
| Treatment | 0.000190 | 2 | 0.6688 | 0.5130784 | |
| Time | 0.000076 | 1 | 0.5386 | 0.4635551 | |
| Replicate:Genotype | 0.000121 | 2 | 0.4275 | 0.6525187 | |
| Replicate:Treatment | 0.001797 | 4 | 3.1689 | 0.0142203 | * |
| Genotype:Treatment | 0.000726 | 2 | 2.5600 | 0.0789358 | . |
| Replicate:Time | 0.002094 | 2 | 7.3828 | 0.0007374 | *** |
| Genotype:Time | 0.000587 | 1 | 4.1384 | 0.0427729 | * |
| Treatment:Time | 0.000432 | 2 | 1.5217 | 0.2199662 | |
| Replicate:Genotype:Treatment | 0.001482 | 4 | 2.6130 | 0.0354782 | * |
| Replicate:Genotype:Time | 0.000435 | 2 | 1.5347 | 0.2171624 | |
| Replicate:Treatment:Time | 0.000228 | 4 | 0.4027 | 0.8066526 | |
| Genotype:Treatment:Time | 0.001148 | 2 | 4.0493 | 0.0183643 | * |
| Replicate:Genotype:Treatment:Time | 0.000396 | 4 | 0.6978 | 0.5939530 | |
| Residuals | 0.043957 | 310 | | | |

---
Signif. codes:  0 ‘***’ 0.001 ‘**’ 0.01 ‘*’ 0.05 ‘.’ 0.1 ‘ ’ 1

#### Figure S5 – 7mth:

| | Sum Sq | Df | F value | Pr(>F) |
|---|---|---|---|---|
| Replicate | 6.3200e-08 | 2 | 0.8794 | 0.4161 |
| Genotype | 3.5900e-08 | 1 | 0.9980 | 0.3186 |
| Treatment | 7.5200e-08 | 2 | 1.0462 | 0.3525 |
| Time | 3.6200e-08 | 1 | 1.0060 | 0.3167 |
| Replicate:Genotype | 5.9000e-08 | 2 | 0.8210 | 0.4409 |
| Replicate:Treatment | 1.3340e-07 | 4 | 0.9275 | 0.4482 |
| Genotype:Treatment | 7.5700e-08 | 2 | 1.0520 | 0.3505 |
| Replicate:Time | 6.3500e-08 | 2 | 0.8833 | 0.4144 |
| Genotype:Time | 3.6300e-08 | 1 | 1.0087 | 0.3160 |
| Treatment:Time | 7.6700e-08 | 2 | 1.0669 | 0.3453 |
| Replicate:Genotype:Treatment | 1.2530e-07 | 4 | 0.8710 | 0.4816 |
| Replicate:Genotype:Time | 5.9500e-08 | 2 | 0.8278 | 0.4380 |
| Replicate:Treatment:Time | 1.3560e-07 | 4 | 0.9429 | 0.4393 |
| Genotype:Treatment:Time | 7.7100e-08 | 2 | 1.0721 | 0.3435 |
| Replicate:Genotype:Treatment:Time | 1.2930e-07 | 4 | 0.8991 | 0.4647 |
| Residuals | 1.1147e-05 | 310 | | |

#### Figure S5 – 7mso:

| | Sum Sq | Df | F value | Pr(>F) | |
|---|---|---|---|---|---|
| Replicate | 0.003994 | 2 | 8.4325 | 0.0002716 | *** |
| Genotype | 0.000310 | 1 | 1.3092 | 0.2534176 | |
| Treatment | 0.003018 | 2 | 6.3717 | 0.0019416 | ** |
| Time | 0.001128 | 1 | 4.7626 | 0.0298355 | * |
| Replicate:Genotype | 0.001380 | 2 | 2.9131 | 0.0557945 | . |
| Replicate:Treatment | 0.009623 | 4 | 10.1569 | 9.721e-08 | *** |
| Genotype:Treatment | 0.001830 | 2 | 3.8627 | 0.0220300 | * |
| Replicate:Time | 0.005042 | 2 | 10.6441 | 3.382e-05 | *** |
| Genotype:Time | 0.000923 | 1 | 3.8990 | 0.0492019 | * |
| Treatment:Time | 0.002408 | 2 | 5.0830 | 0.0067286 | ** |
| Replicate:Genotype:Treatment | 0.000517 | 4 | 0.5455 | 0.7024501 | |
| Replicate:Genotype:Time | 0.000427 | 2 | 0.9011 | 0.4071687 | |
| Replicate:Treatment:Time | 0.002878 | 4 | 3.0376 | 0.0176781 | * |
| Genotype:Treatment:Time | 0.001822 | 2 | 3.8463 | 0.0223856 | * |
| Replicate:Genotype:Treatment:Time | 0.001029 | 4 | 1.0862 | 0.3633857 | |
| Residuals | 0.073424 | 310 | | | |

---
Signif. codes: 0 '***' 0.001 '**' 0.01 '*' 0.05 '.' 0.1 ' ' 1

#### Figure S5 – 8mto:

| | Sum Sq | Df | F value | Pr(>F) | |
|---|---|---|---|---|---|
| Replicate | 0.001923 | 2 | 8.9858 | 0.0001609 | *** |
| Genotype | 0.001366 | 1 | 12.7649 | 0.0004094 | *** |
| Treatment | 0.000404 | 2 | 1.8857 | 0.1534555 | |
| Time | 0.000155 | 1 | 1.4510 | 0.2292870 | |
| Replicate:Genotype | 0.001268 | 2 | 5.9248 | 0.0029845 | ** |
| Replicate:Treatment | 0.001073 | 4 | 2.5062 | 0.0421856 | * |
| Genotype:Treatment | 0.000914 | 2 | 4.2710 | 0.0147993 | * |
| Replicate:Time | 0.000342 | 2 | 1.5990 | 0.2037695 | |
| Genotype:Time | 0.000024 | 1 | 0.2239 | 0.6364433 | |
| Treatment:Time | 0.000304 | 2 | 1.4182 | 0.2437204 | |
| Replicate:Genotype:Treatment | 0.000655 | 4 | 1.5304 | 0.1931583 | |
| Replicate:Genotype:Time | 0.000079 | 2 | 0.3706 | 0.6906143 | |
| Replicate:Treatment:Time | 0.000627 | 4 | 1.4648 | 0.2127574 | |
| Genotype:Treatment:Time | 0.000412 | 2 | 1.9244 | 0.1477047 | |
| Replicate:Genotype:Treatment:Time | 0.001212 | 4 | 2.8308 | 0.0248535 | * |
| Residuals | 0.033172 | 310 | | | |

---
Signif. codes: 0 '***' 0.001 '**' 0.01 '*' 0.05 '.' 0.1 ' ' 1

#### Figure S5 – 8mso:

| | Sum Sq | Df | F value | Pr(>F) | |
|---|---|---|---|---|---|
| Replicate | 0.25250 | 2 | 21.7777 | 1.413e-09 | *** |
| Genotype | 0.15321 | 1 | 26.4280 | 4.854e-07 | *** |
| Treatment | 0.06518 | 2 | 5.6220 | 0.0039962 | ** |
| Time | 0.01381 | 1 | 2.3821 | 0.1237491 | |
| Replicate:Genotype | 0.02493 | 2 | 2.1502 | 0.1181932 | |
| Replicate:Treatment | 0.11608 | 4 | 5.0059 | 0.0006365 | *** |
| Genotype:Treatment | 0.02660 | 2 | 2.2943 | 0.1025428 | |
| Replicate:Time | 0.06951 | 2 | 5.9950 | 0.0027892 | ** |
| Genotype:Time | 0.04122 | 1 | 7.1111 | 0.0080629 | ** |
| Treatment:Time | 0.03398 | 2 | 2.9309 | 0.0548299 | . |
| Replicate:Genotype:Treatment | 0.02968 | 4 | 1.2801 | 0.2776854 | |
| Replicate:Genotype:Time | 0.04329 | 2 | 3.7334 | 0.0249937 | * |
| Replicate:Treatment:Time | 0.03396 | 4 | 1.4645 | 0.2128476 | |
| Genotype:Treatment:Time | 0.03440 | 2 | 2.9671 | 0.0529165 | . |
| Replicate:Genotype:Treatment:Time | 0.01630 | 4 | 0.7031 | 0.5903061 | |
| Residuals | 1.79713 | 310 | | | |

---
Signif. codes: 0 '***' 0.001 '**' 0.01 '*' 0.05 '.' 0.1 ' ' 1
